# Mechanistic origin of different binding affinities of SARS-CoV and SARS-CoV-2 spike RBDs to human ACE2

**DOI:** 10.1101/2022.02.05.479221

**Authors:** Zhi-Bi Zhang, Yuan-Ling Xia, Jian-Xin Shen, Wen-Wen Du, Yun-Xin Fu, Shu-Qun Liu

## Abstract

The receptor-binding domain (RBD) of the SARS-CoV-2 spike protein mediates viral entry into host cells through binding to the cell-surface receptor angiotensin-converting enzyme 2 (ACE2). It has been shown that SARS-CoV-2 RBD (RBD_CoV2_) has a higher binding affinity to human ACE2 than its highly homologous SARS-CoV RBD (RBD_CoV_), for which the mechanistic reasons still remain to be elucidated. Here, we used the multiple-replica molecular dynamics (MD) simulations, molecular mechanics Poisson-Boltzmann surface area (MM-PBSA) binding free energy calculations, and interface residue contact network (IRCN) analysis approach to explore the mechanistic origin of different ACE2 binding affinities of these two RBDs. The results demonstrate that, when compared to the RBD_CoV2_-ACE2 complex, the RBD_CoV_-ACE2 complex features the enhanced overall structural fluctuations and inter-protein positional movements and increased conformational entropy and diversity. The inter-protein electrostatic attractive interactions are a dominant force in determining the high ACE2 affinities of both RBDs, while the significantly strengthened electrostatic forces of attraction of ACE2 to RBD_CoV2_ determine the higher ACE2 binding affinity of RBD_CoV2_ than of RBD_CoV_. Comprehensive comparative analyses of the residue binding free energy components and IRCNs reveal that, although any RBD residue substitution involved in the charge change can significantly impact the inter-protein electrostatic interaction strength, it is the substitutions at the RBD interface that lead to the overall stronger electrostatic attractive force of RBD_CoV2_-ACE2, which in turn not only tightens the interface packing and suppresses the dynamics of RBD_CoV2_-ACE2, but also enhances the ACE2 binding affinity of RBD_CoV2_ compared to that of RBD_CoV_. Since the RBD residue substitutions involving gain/loss of the positively/negatively charged residues, in particular those near/at the binding interfaces with the potential to form hydrogen bonds and/or salt bridges with ACE2, can greatly enhance the ACE2 binding affinity, the SARS-CoV-2 variants carrying such mutations should be paid special attention to.

## 1. Introduction

The severe acute respiratory syndrome coronavirus 2 (SARS-CoV-2) [1,2], which shares high homology with the SARS-CoV (about 80% nucleotide sequence identical at the genome level) responsible for the SARS outbreak in 2002-2003 [3], causes the ongoing coronavirus disease 2019 (COVID-19). As of January 2022, SARS-CoV-2 has caused over 340 million cases of COVID-19 and more than 5.5 million deaths worldwide (https://www.who.int/emergencies/diseases/novel-coronavirus-2019 (accessed on 24 January 2022)).

Both SARS-CoV and SARS-CoV-2 belong to the *Sarbecovirus* subgenus of betacoronaviruses [4] and utilize the same receptor, angiotensin-converting enzyme 2 (ACE2) for cell entry, an infection process triggered by attachment of the virus spike protein to the cell-surface receptor [5,6]. The spike protein of coronaviruses (CoV) is a class I membrane fusion glycoprotein [7] composed of three identical chains/protomers (hence the term homotrimeric spike or spike trimer) protruding from the surface of lipid-enveloped CoV particles [8,9]. Each protomer consists of two subunits, S1 and S2, which are post-translationally cleaved products from the single-chain polypeptide spike precursor and are responsible for virus attachment to cells and fusion of the viral and cellular membranes, respectively [7,10-12]. In the spike trimer structure, N- and C-terminal portions of each S1 subunit fold independently as two large domains, the N-terminal domain (NTD) and the C-terminal domain (CTD), with the latter serving as the receptor-binding domain (RBD) in the cases of SARS-CoV and SARS-CoV-2 [13-18]. The S1 subunit, especially the RBD on it, is also the immunodominant target of the humoral response and hence the uppermost component of both mRNA and adenovirus-based vaccines [19,20].

The spike trimer on the virus surface does not present as a single rigid conformation but rather as an ensemble of different conformations/states (i.e., the closed state and the open states with one, two, or three erect RBD(s)) that coexist in equilibrium with different population distributions [17,18,21]. In the closed conformation, all the three RBDs lie down and pack tightly against one another and the NTD of the anticlockwise neighboring protomer, thus capping the top of the central helical stalk of S2 subunits and simultaneously leaving the ACE2-binding surface (i.e., partial surface of the receptor binding motif (RBM) that interacts directly with ACE2 [17] buried inside the trimer and inaccessible to ACE2. In the absence of ACE2, the closed state of the spike trimer can spontaneously convert to the open states through a hinge-like motion that progressively raises RBDs, thus allowing the recognition and binding to ACE2 due to the full exposure of RBM on the erect RBDs [22]. Upon binding, the first ACE2-bound RBD is stabilized in the open orientation, which facilitates the other two RBDs to consecutively stand up followed by binding to ACE2 until a fully open, three-ACE2-bound structure is formed that primes the spike trimer for S2 unsheathing required for membrane fusion [18].

Although the erection of RBD is a prerequisite for its binding to ACE2, RBD is an independently folded domain for which the erection has only marginal impact on its overall conformation [18]. As a result, the binding affinity of the spike protein to ACE2 was usually evaluated using RBD rather than the spike trimer. Interestingly, multiple earlier experimental and simulation studies have collectively demonstrated that the SARS-CoV-2 RBD (RBD_CoV2_) has higher binding affinity to ACE2 than SARS-CoV RBD (RBD_CoV_) [17,23-28], and this likely explains why SARS-CoV-2 manifests increased infectivity and transmissibility in comparison to SARS-CoV.

The crystal structures of RBD_CoV_ and RBD_CoV2_ in complex with human ACE2 have been determined at high resolution [13,14], revealing that the two RBDs share not only overall similar conformations (with C_α_ root mean square deviation (RMSD) of 0.47 Å) but also nearly identical modes of binding to ACE2 (Figures 1A to C). Both RBDs have two subdomains: a core that is formed by a twisted five-stranded antiparallel β-sheet connected by short helices and loops, and an extended insertion of RBM composed of a short two-stranded antiparallel β-sheet, two short helices, and several long loops. Although the core contains few residues capable of making contacts with ACE2, most of the ACE2-contacting residues are from RBM. The sequence identity shared by the RBD_CoV_ and RBD_CoV2_ is 73.2%, which explains why the two RBDs have highly similar overall structures. However, the sequence identity of cores increases to 88.0% and that of RBMs falls to 47.8%, and this may be related to different ACE2 binding affinities of the two RBDs due to more residue changes and much more ACE2-contacting residues in RBM than in the core (Figure 1 D).

**Figure 1.**
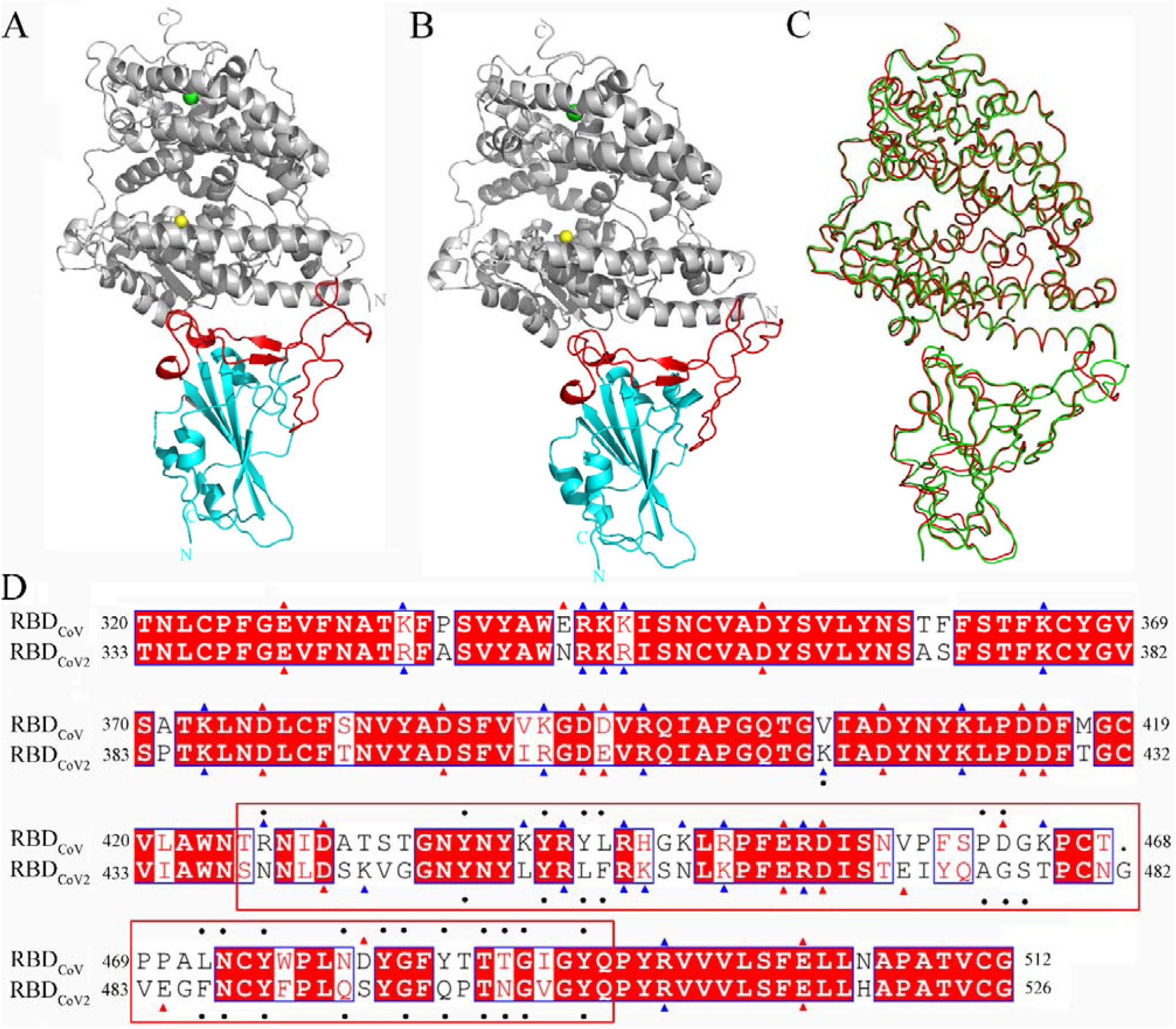
Structures of the RBD_CoV_-ACE2 and RBD_CoV2_-ACE2 complexes and sequence alignment of RBD_CoV_ and RBD_CoV2_. (A) and (B) Cartoon representations of the complete complex structures of RBD_CoV_-ACE2 (modeled based on the crystal structure with PDB ID 2AJF [13]) and RBD_CoV2_-ACE2 (PDB ID: 6M0J [14]), respectively. ACE2 is colored gray, with Zn^2+^ and Cl^−^ ions represented as spheres in yellow and green, respectively; the cores and RBMs of both RBDs are colored cyan and red, respectively. (C) Backbone superposition of the complex structures of RBD_CoV_-ACE2 (red) and RBD_CoV2_-ACE2 (green). (D) Structure-based sequence alignment of RBD_CoV_ and RBD_CoV2_. The identical residues are white on red background and the similar residues are red on white background; the negatively and positively charged residues are indicated by red and blue triangles, respectively; the ACE2-contacting residues (or RBD interface residues) identified in this work are indicated by black dots; RBM (residues 438-506 according to residue numbering of RBD_CoV2_) is highlighted by enclosure with a red box.

Although multiple previous studies have provided insights into the structural and molecular basis responsible for the different binding affinities of RBD_CoV_ and RBD_CoV2_ to human ACE2 [23,24,29-32], the underlying mechanisms modulating the mechanics and energetics of RBD-ACE2 interactions still remain to be elucidated. In order to explore the mechanistic reasons for the experimentally observed difference in ACE2 binding affinities of the two RBDs, in this work we performed multiple-replica MD simulations on the structures of the RBD_CoV_-ACE2 and RBD_CoV2_-ACE2 complexes, followed by comparative analyses in terms of dynamics and thermodynamics, calculations of the protein-protein and per-residue binding free energies, constructions of the interface residue contact networks (IRCNs), and comprehensive comparative analyses of IRCNs, interface interactions, and binding free energy components of individual residues. The results presented in this paper shed light on the dynamic and energetic aspects of the modulation mechanism of RBD-ACE2 interactions, explain why RBD_CoV2_ enhances its binding affinity to ACE2 compared to that of RBD_CoV_, and may also help surveil the emergence of novel SARS-CoV-2 variants.

## 2. Materials and Methods

### 2.1. Structural preparation

The X-ray crystallographic structures of the RBD_CoV_-ACE2 and RBD_CoV2_-ACE2 complexes were obtained from Protein Data Bank (PDB; http://www.rcsb.org (accessed on 8 August 2021)) with PDB IDs of 2AJF [13] and 6M0J [14], respectively. The hetero atoms and crystallographic water molecules included in the PDB files were removed while the atomic coordinates for ACE2, RBD, and ACE2-bound Zn^2+^ and Cl^−^ ions were retained. For the RBD_CoV_-ACE2 complex, the missing atomic coordinates of the RBD_CoV_ residues 320-322, 376-381 and 503-512 were modeled using the SWISS-MODEL server [33] with the RBD_CoV2_ structure as the template. The structure-based sequence alignment between RBD_CoV_ and RBD_CoV2_ was performed using the Dali server (http://ekhidna.biocenter.helsinki.fi/dali/ (accessed on 12 August 2021)) [34] and visualized by ESPript 3.0 (https://espript.ibcp.fr/ESPript/cgi-bin/ESPript.cgi (accessed on 12 August 2021)) [35]. The RBD residue numbering is according to that of RBD_CoV2_ unless otherwise indicated.

### 2.2 MD simulations

All MD simulations were performed using the GROMACS 5.1.4 software package [36] with AMBER99SB-ILDN force field [37]. The p*K*_a_ values of all titratable residues in both complex structures were calculated using the PropKa server (https://www.ddl.unimi.it/vegaol/propka.htm (accessed on 12 August 2021)) [38] to assign their protonation states at pH 7.4. Since the side-chain p*K*_a_ values of all lysines and arginines and all glutamate and aspartate residues are greater and less than 7.4, respectively, they were protonated (positively charged) and deprotonated (negatively charged), respectively; since His374 of ACE2 in both complexes and His493 of ACE2 in RBD_CoV2_-ACE2 have predicted side-chain p*K*_a_ values greater than 7.4, their imidazole N_δ1_ and N_ε2_ atoms were both protonated (positively charged); all the other histidines have side-chain p*K*_a_ values less than 7.4 and hence were treated as uncharged neutral residues, with the imidazole rings being protonated automatically on either N_δ1_ or N_ε2_ based on the optimal hydrogen-bonding conformation using the GROMACS tool ‘gmx pdb2gmx’.

After protonation, each complex structure was solvated with the TIP3P water model [39] in a periodic dodecahedron box with a minimum solute-box wall distance of 1.2 nm. The net charges of both systems were neutralized with NaCl at the physiological concentration of 0.15 M. Each system was subjected to a steepest descent energy minimization until no significant energy change could be detected. To effectively ‘soak’ the solute into the solvent, four consecutive 100-ps NVT MD simulations were conducted at 310 K, with the protein heavy atoms restrained by decreasing harmonic potential force constants of 1000, 100, 10, and 0 kJ/mol/nm^2^. To improve the sampling of the complex conformational space, each system was subjected to 10 independent 15-ns production MD simulations, with each replica initialized with different initial atomic velocities assigned from a Maxwell distribution at 310 K. In the production MD runs, the LINear Constraint Solver (LINCS) algorithm [40] was used to constrain all bonds to equilibrium lengths, thus allowing a time step of 2 fs; the particle-mesh Ewald (PME) method [41] was used to treat the long-range electrostatic interactions with a real-space cut-off of 1.0 nm, grid spacing of 0.12 nm, and interpolation order of 4; van der Waals (vdW) interactions were treated using the Verlet scheme with a cut-off distance of 1.0 nm; the system temperature was controlled at 310 K using the velocity-rescaling thermostat [42] with a time constant of 0.1 ps; the system pressure was maintained at 1 atm using the Parrinello-Rahman barostat [43] with a time constant of 2.0 ps; and system coordinates were saved every 2 ps.

### 2.3 MD trajectory analysis

For the obtained production MD trajectories of the 10 replicas, the time-dependent C_α_ RMSD values relative to the starting structure were calculated using the GROMACS tools ‘gmx rms’. For each complex system, the equilibrated portion of each of the 10 replicas was concatenated into a single joined equilibrium trajectory, based on which the principal component analysis (PCA) was performed on the C_α_ atoms using the GROMACS tools ‘gmx covar’ and ‘gmx anaeig’. Then, the first two eigenvector projections were used as the reaction coordinates to reconstruct the two-dimensional free energy landscape (FEL) [44-46] with the following equation:

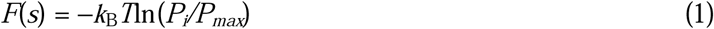

where *k*_B_ is Boltzmann’s constant, *T* is the simulation temperature, *P*_*i*_ is the probability of finding the system in state *i* characterized by the two reaction coordinates (i.e., the first two eigenvectors), and *P*_*max*_ is the probability of the most probable state.

For each complex, 100 conformations were randomly extracted from the global free energy minimum of FEL, and were treated as the representative structures for the subsequent binding free energy calculation and interaction/contact network analysis.

### 2.4 Binding free energy calculation

The binding free energies of RBD_CoV_ and RBD_CoV2_ to ACE2 were evaluated using the molecular mechanics Possion-Boltzmann surface area (MM-PBSA) method as implemented in the GROMACS tool g_mmpbsa [47]. MM-PBSA is an endpoint approach capable of estimating the protein-protein/ligand binding free energy based merely on the structure or structural ensemble of the bound complex without considering either the physical or the non-physical intermediates [48]. Here the binding free energy was estimated with the single trajectory approach, which assumes that the conformations of the free RBD and free ACE2 are identical to those in the protein-protein complex.

In MM-PBSA, the binding free energy (Δ*G*_*binding*_) of a single protein-protein complex was calculated as following:

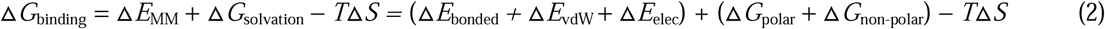

where Δ*E*_*MM*_, Δ*G*_*solvation*_, and *T*Δ*S* are the change in the vacuum molecular mechanics potential energy, the change in the solvation free energy, and the solute entropy change, respectively, upon the formation of the complex from the two free protein partners; Δ*E*_*MM*_ is further decomposed into the bonded energy change (Δ*E*_*bonded*_) and the changes in the van der Waals (Δ*E*_*vdW*_) and electrostatic (Δ*E*_*elec*_) potential energies, with Δ*E*_*bonded*_ value being zero due to the single trajectory approach, and Δ*E*_*vdW*_ and Δ*E*_*elec*_ being the vdW and electrostatic interaction energies between the two binding partners, respectively; and Δ*G*_*solvation*_ is decomposed into the polar (Δ*G*_*polar*_) and non-polar (Δ*G*_*non-polar*_) solvation free energy contributions, with Δ*G*_*polar*_ being actually the electrostatic desolvation free energy upon binding and Δ*G*_*non-polar*_ representing the hydrophobic effect arising from the solvent entropy gain upon binding [48-50]. The term *T*Δ*S* was not included in our calculations because i) estimating the solute entropy change is a time-consuming task while often producing highly uncertain results with a larger standard error than those of the other energy terms [51-53] and ii) omitting the solute entropy term likely has only minor impact on comparison between the relative binding free energies of different ligands/proteins to the same receptor protein [47,51].

For each complex, MM-PBSA calculations were performed on the structural ensemble of the 100 representative conformations. The g_mmpbsa default parameters were used with the exceptions of those for calculating the polar solvation free energy; specifically, *G*_*polar*_ was calculated by solving the nonlinear PB equation (npbe) with grid resolution (gridspace) of 0.4 Å, ionic strength (NaCl concentration) of 0.15 M, and solute (pdie) and solvent (sdie) dielectric constants of 4 and 80, respectively.

Per-residue contribution to the overall binding free energy (hereafter referred to as the residue binding free energy) of each complex was obtained by implementing the ‘binding energy decomposition’ module of g_mmpbsa, which allows to decompose the total calculated binding energy into contributions from individual residues by calculating each atom’s energy components (i.e., *E*_*MM*_, *G*_*polar*_, and *G*_*nonpolar*_) of a residue in both the free and bound forms.

### 2.5 Interface interaction/contact analyses

The interface residue contact network (IRCN) was constructed based on the numbers of close inter-atomic contacts of involved residues from RBD and ACE2. A close inter-atomic contact is considered to exist if the “overlap” value between any two atoms is greater than −4.0 Å. The overlap between atoms *i* and *j* (*overlap*_*ij*_) is defined as:

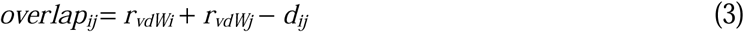

where *r*_*vdWi*_ and *r*_*vdWj*_ represent the vdW radii of atoms *i* and *j*, respectively, and *d*_*ij*_ denotes the distance between the nuclei of them. A hydrogen bond (HB) is considered to exist if the donor-acceptor distance is less than 3.5 Å and the donor-hydrogen-acceptor angle is greater than 120°. A salt bridge (SB) is considered to be formed if the distance between any side-chain oxygen atom of a negatively charged residue (Asp or Glu) and any side-chain nitrogen atom of a positively charged residue (Arg or Lys) is less than 4.0 Å.

For each representative structure of the RBD_CoV_-ACE2 and RBD_CoV2_-ACE2 complexes, inter-atomic close contacts, HBs, and SBs were identified using the Chimera ‘Find Contacts’ tool [54] and VMD plugins ‘Hydrogen Bonds’ and ‘Salt Bridge’ [55], respectively. The average number of close inter-atomic contacts in a specific contacting residue pair was calculated over the 100 representative structures; only the residues with the average contact number greater than 1.0 were considered as the binding interface residues. The occupancy of a HB or SB was calculated as the fraction of the structures within which a specified HB/SB exists out of the 100 representative structures; only those with occupancy greater than 20% were considered as stable HBs/SBs. Finally, the IRCN and interface HB were generated from the 100 representative structures of each complex by Chimera 1.14 [54] and visualized by Cytoscape 3.8.1 [56]. The distance of a given residue to the binding interfaces, which is defined as the minimum distance between any pair of atoms from the given residue and from the interface residues, was calculated over the 100 representative structures using the GROMACS tools ‘gmx pairdist’.

## 3. Results

### 3.1 Structural fluctuations during simulations

Figure 2 shows the time-dependent C_α_ RMSD values of the two complexes relative to their respective starting structures during the multiple-replica simulations. All the 10 replicas of RBD_CoV2_-ACE2 require only a few ps to reach relatively stable RMSD values (Figure 2B), whereas some replicas of RBD_CoV_-ACE2 require more than 1.3 ns to reach a relative plateau of RMSD values (Figure 2A). In order to ensure the data temporal consistency of the two simulation systems, we arbitrarily treated the 2-15-ns trajectory of each replica as the equilibrated portion. It is clear that i) the equilibrated portions of the 10 RBD_CoV_-ACE2 replicas span a wider RMSD range (0.14-0.49 nm) than those of the 10 RBD_CoV2_-ACE2 replicas (0.11-0.31 nm) and ii) RBD_CoV_-ACE2 has more replicas with the equilibrium fluctuation amplitude greater than 0.15 nm than RBD_CoV2_-ACE2, indicating that during the equilibrium the RBD_CoV_-ACE2 complex not only deviated more from its starting conformation but also experienced larger global structural fluctuations than did the RBD_CoV2_-ACE2 complex. Nevertheless, visual inspection of all replica trajectories revealed that both RBD_CoV_ and RBD_CoV2_ bound stably to ACE2 throughout the 15-ns simulations.

**Figure 2.**
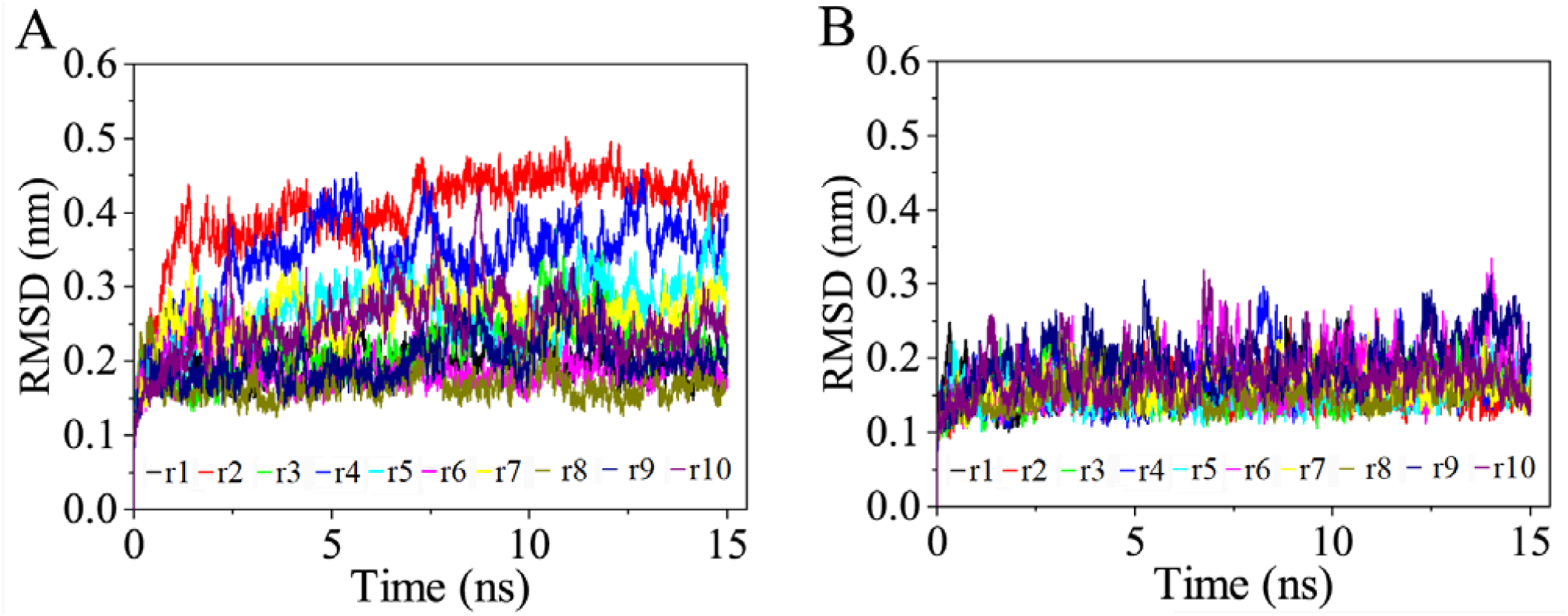
Time evolution of the C_α_ root mean square deviation (RMSD) values of the two complexes relative to their respective starting structures calculated from the 10 independent MD simulation replicas (r1-10). (A) RBD_CoV_-ACE2 complex. (B) RBD_CoV2_-ACE2 complex.

In order to further evaluate the structural fluctuations of the two binding partners and their relative mobility with respect to each other, we calculated the time-dependent C_α_ RMSD values of the RBD and ACE2 after least squares fitting to either the respective structure (self-fitting) or to the structure of the other partner (non-self-fitting) in the starting complex. The results (Figure S1) reveal that, for both complexes, the self-fitting RMSD values of RBDs (Figures S1A and B) and ACE2s (Figures S1C and D) are clearly lower than their respective non-self-fitting values (Figures S1E-H). This is not surprising, as the self-fitting RMSD values provide information about the intrinsic fluctuations of a bound protein in the complex, while non-self-fitting RMSD values mainly reflect the relative movements of one protein with respect to the other in the complex. The self-fitting RMSD curves of RBD and ACE2 span a wider width and have a larger fluctuation amplitude in the RBD_CoV_-ACE2 (Figures S1A and C) than in the RBD_CoV2_-ACE2 complex (Figures S1B and D), respectively, implying that the two binding partners have a larger structural variability in the former than in the latter complex. Interestingly, for both complexes, although their non-self-fitting RMSD curves of RBD and ACE2 collectively show drastic fluctuations, the obviously wider RMSD curve widths observed for the RBD_CoV_-ACE2 complex indicate that its two binding partners experienced larger relative position shifts/movements than did the partners in the RBD_CoV2_-ACE2 complex.

Taken together, when compared to the RBD_CoV2_-ACE2 complex, the RBD_CoV_-ACE2 complex experienced not only more drastic structural fluctuations at both the levels of the entire complex and individual bound proteins, but also larger relative movements between the two bound proteins. Thus, the RBD_CoV_-ACE2 complex has a lower overall structural stability, larger structural variabilities of the two bound proteins, and a less stable binding orientation between the two partners than the RBD_CoV2_-ACE2 complex.

### 3.2 Principal components and free energy landscapes

PCA analysis was performed on the single joined equilibrium trajectory of each complex to extract the most important PCs/eigenvectors. For the RBD_CoV_-ACE2 and RBD_CoV2_-ACE2 complexes, the first two eigenvectors possess the largest eigenvalues (Figure S2), and their cumulative eigenvalues contribute 62.0% and 48.5% to the total mean square fluctuation values of the two complexes, respectively (Figure S2, inset). Since the conformational space is spanned by 3N (N is the number of C_α_ atoms; 790 and 791 for RBD_CoV_-ACE2 and RBD_CoV2_-ACE2, respectively) eigenvectors, it is reasonable to believe that first two eigenvectors contribute substantially to the overall conformational freedom in the space and, therefore, the essential subspace spanned by the first two eigenvectors contains the main conformational states/substates sampled by the MD simulations, from which the most representative stable conformations can be identified through reconstructing the FEL.

Figure 3 shows the two-dimensional FELs of the two complexes using the first two eigenvector projections as the reaction coordinates. It is clear that the FEL of RBD_CoV_-ACE2 (Figure 3A) covers a larger region in the essential subspace than that of RBD_CoV2_-ACE2 (Figure 3B), indicating a larger conformational entropy of the former than of the latter complex. Furthermore, the FEL of RBD_CoV_-ACE2 features a rough/rugged surface because it contains two large basins (with a free energy level lower than −10 kJ/mol) within which multiple local minima (with a free energy level lower than −11 or −12 kJ/mol) are present, and that of RBD_CoV2_-ACE2 shows a typical funnel-like shape characterized by a continuous falling of free energy until reaching the global free energy minimum (−14 kJ/mol) but without local minima observed on the funnel wall. Therefore, it can be considered that during simulations the RBD_CoV_-ACE2 and RBD_CoV2_-ACE2 complexes sampled two and one conformational states, respectively, with the former’s two states containing multiple metastable substates while the latter’s single state in the global minimum being its most stable state. Nevertheless, the global free energy minimum within one of the two basins of RBD_CoV_-ACE2’s FEL has an equivalent free energy level (−14 kJ/mol) to that of RBD_CoV2_-ACE2’s global minimum, indicating the equivalent thermostability of the two complexes. Interestingly, the global minimum of RBD_CoV_-ACE2’s FEL has a smaller size than that of RBD_CoV2_-ACE2’s FEL, indicating that the latter complex sampled a larger population of the most thermostable conformations during the MD simulations.

**Figure 3.**
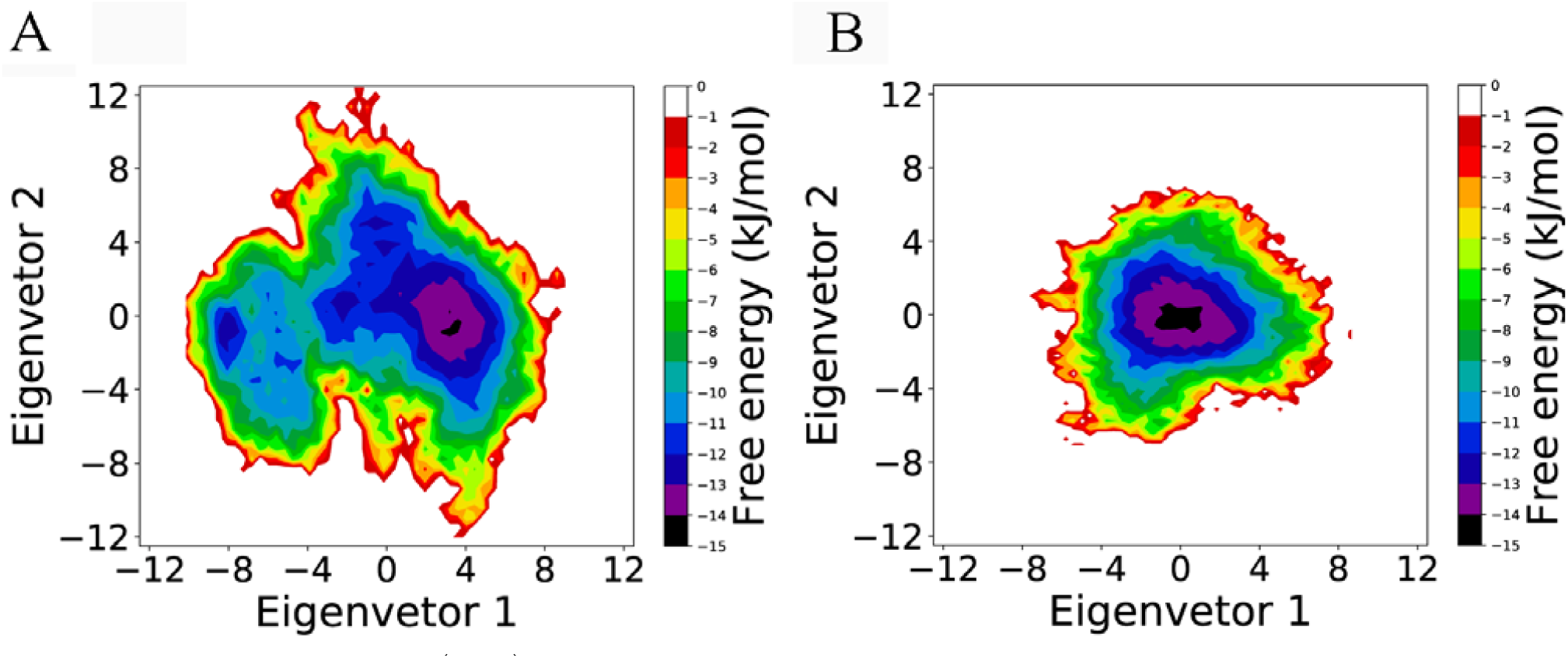
Free energy landscapes (FELs) of the two complexes as a function of the projection of the joined equilibrium trajectory onto the essential subspace spanned by eigenvectors 1 and 2. (A) FEL of the RBD_CoV_-ACE2 complex. (B) FEL of the RBD_CoV2_-ACE2 complex. The color bar represents the relative free energy value in kJ/mol.

In summary, although the RBD_CoV_-ACE2 complex has a larger conformational entropy and richer conformational diversity than RBD_CoV2_-ACE2, these two complexes present the equivalent thermostability. Therefore, it is reasonable to take the most thermostable conformations as the representative structures of the two complexes. As a result, 100 conformations/structures were randomly extracted from the global free energy minimum of each FEL for the subsequent binding free energy calculation and interface contact network analysis.

### 3.4 Binding free energy calculation

Table 1 shows the average values and corresponding standard deviations (SDs) of binding free energy components for the two complexes. The calculated average values of the binding free energy (ΔG_*binding*_) for RBD_CoV_-ACE2 and RBD_CoV2_-ACE2 are −2289.2 and −2455.0 kJ/mol, respectively; although the ranges of Δ*G*_*binding*_ values of the two complexes will become overlapping when taking SDs into account, the p-value of 2.1 × 10^−19^ by the one-sided t-test indicates that the binding free energy of RBD_CoV2_-ACE2 is statistically significantly lower than that of RBD_CoV_-ACE2, consistent with previous experimental observations demonstrating that RBD_CoV2_ has a higher binding affinity than RBD_CoV_ to human ACE2 receptor [17,23,27,28].

**Table 1.** Average values (SDs are in parentheses) of various MM-PBSA energy terms (kJ/mol) of the RBD_CoV_-ACE2 and RBD_CoV2_-ACE2 complexes calculated over their respective 100 representative structures.

| energy term<sup>a</sup> | RBD<sub>CoV</sub>-ACE2 | RBD<sub>CoV2</sub>-ACE2 |
| --- | --- | --- |
| $\Delta E_{\text{vdW}}$ | −360.2 (27.8) | −376.0 (24.0) |
| $\Delta E_{\text{elec}}$ | −2338.4 (146.8) | −2597.0 (146.7) |
| $\Delta G_{\text{polar}}$ | 452.3 (85.5) | 561.9 (84.8) |
| $\Delta G_{\text{non-polar}}$ | −42.9 (4.2) | −43.9 (3.4) |
| $\Delta G_{\text{binding}}$ | −2289.2 (129.2) | −2455.0 (103.2) |

<sup>a</sup> For detailed explanations of the energy terms, see Equation 2 in the section 2.4.

It is clear that for each complex, the electrostatic interaction potential energy (Δ*E*_elec_) contributes most significantly to lowering the binding free energy so that this term alone can over-compensate for the large negative contribution from the electrostatic desolvation energy term (Δ*G*_polar_), whereas the sum of the positive contributions from the vdW interaction potential energy (Δ*E*_vdW_) and solvent exclusion effect (Δ*G*_non-polar_) cannot offset the unfavorable electrostatic desolvation penalty. The difference in the average values of Δ*G*_binding_ from RBD_CoV_-ACE2 to RBD_CoV2_-ACE2 is −165.8 kJ/mol, and the differences in the average values of Δ*E*_vdW_, Δ*E*_elec_, Δ*G*_polar_, and Δ*G*_non-polar_ are −15.9, −258.6, 109.6, and −1.0 kJ/mol, respectively, clearly indicating that the higher binding affinity of RBD_CoV2_ to ACE2 primarily originates from the considerably stronger inter-protein electrostatic attractive interactions in RBD_CoV2_-ACE2 than in RBD_CoV_-ACE2, although the inter-protein vdW interactions and the hydrophobic effect (Δ*G*_non-polar_) are also slightly more favorable in RBD_CoV2_-ACE2 than in RBD_CoV_-ACE2. Interestingly, all the energy terms have larger SDs for RBD_CoV_-ACE2 than for RBD_CoV2_-ACE2, meaning a larger dispersion around their respective average values and hence less tight inter-protein association in the RBD_CoV_-ACE2 complex, in consistence with earlier comparative analysis in terms of RMSD.

Figure 4A shows the ACE2 residues that have the average binding free energy values greater than 20.0 or lower than −20.0 kJ/mol in both complexes. All these residues are charged ones, with the positively charged and negatively charged residues making the negative and positive contributions, respectively, to the binding affinity of ACE2 to both RBDs; furthermore, the magnitudes of the energy values depend on the residue distances to the binding interfaces, i.e., the residues located far from the binding interfaces (marked by light or lighter gray rectangles) generally have a smaller magnitude of the absolute energy values (lower than 40 kJ/mol) than those located near/at the interfaces (greater than 40 kJ/mol), and the large differences in the energy values between the complexes mainly occur on the residues located at/near the interfaces (marked by black or dark gray rectangles). Figure 4C shows the per-residue binding free energy difference from the RBD_CoV_-bound to RBD_CoV2_-bound ACE2s. All the residues with the significant energy differences (greater than 20 or lower than −20 kJ/mol) are charged ones located near/at the binding interfaces with the exception of H493, which is very far from the interfaces (greater than 3.0 nm; see Figure 4A) and has a net charge of 0 and +1 in the RBD_CoV_-bound and RBD_CoV2_-bound ACE2s (for details; see the section 2.2), respectively. Despite its remote distance from the interfaces, the change in the charge property of His493 still leads to a considerable difference in its binding free energy, indicating the importance of the long-range electrostatic interactions in affecting residue’s contribution to the binding affinity. For the charged residues located near/at the interfaces, the large changes in the binding free energy also arise from the differences in the electrostatic interactions, e.g., D30 contributes to enhancing ACE2’s affinity to RBD_CoV2_ due to its stronger electrostatic attractive interactions with RBD_CoV2_ (−226.5 kJ/mol) than with RBD_CoV_ (−66.5 kJ/mol), while K353 contributes to reducing ACE2’s affinity to RBD_CoV2_ due to its stronger electrostatic repulsive interactions with RBD_CoV2_ (46.6 kJ/mol) than with RBD_CoV_ (9.4 kJ/mol; see Table S1).

**Figure 4.**
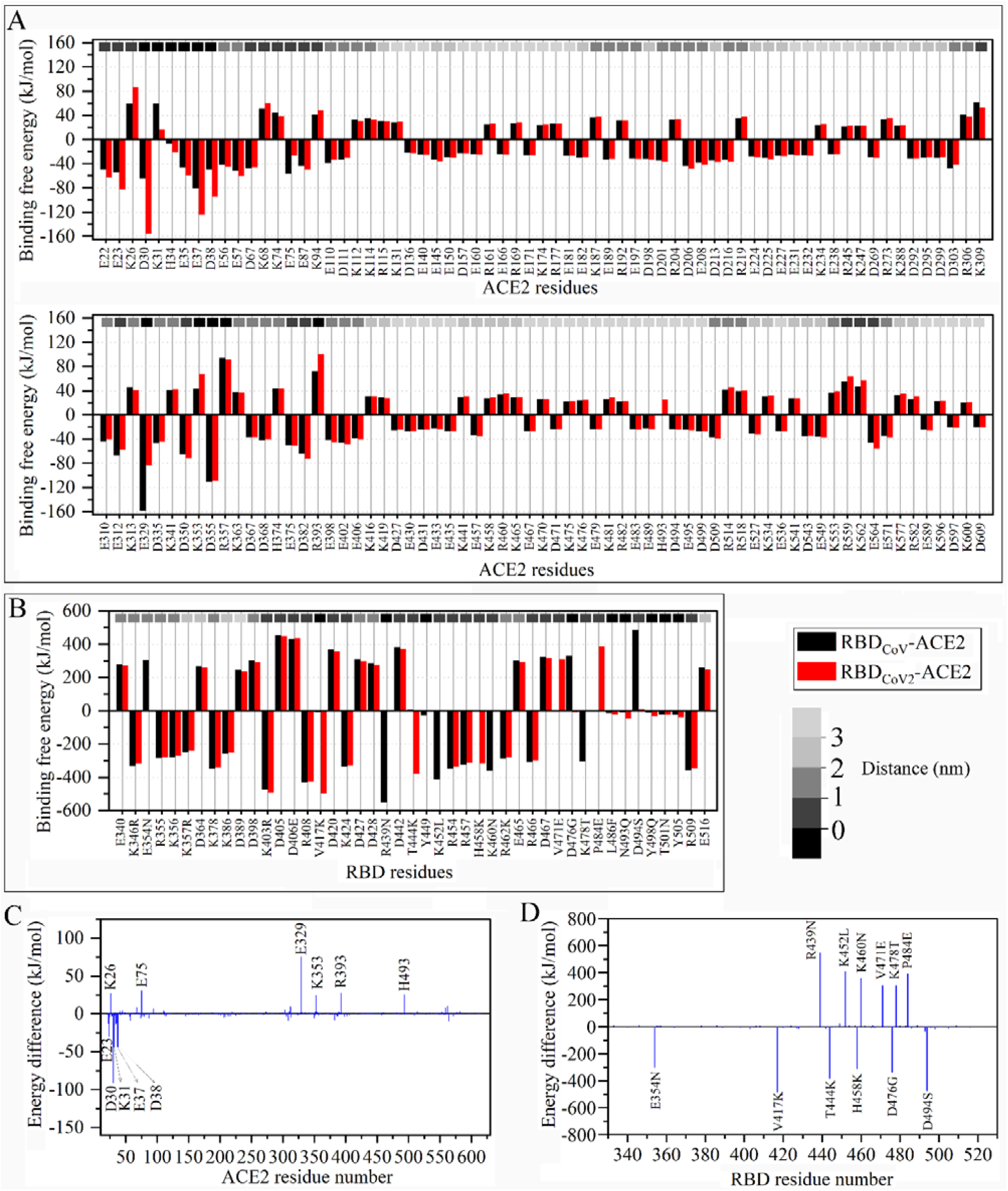
Residue binding free energy average values and their differences between the RBD_CoV_-ACE2 and RBD_CoV2_-ACE2 complexes. (A) Residue binding free energy average values of the RBD_CoV_-bound (black) and RBD_CoV2_-bound (red) ACE2s. (B) Residue binding free energy average values of RBD_CoV_ (black) and RBD_CoV2_ (red). In (A) and (B), only the residues with the energy values greater than 20 kJ/mol or lower than −20 kJ/mol are shown; the distances of residues to the binding interfaces are marked by small rectangles of different shades of gray along the top horizontal axis, with the black rectangles representing the distance of 0 nm (i.e., the interface residues identified in this work) and those of reduced gray representing the increased distance to the binding interfaces as indicated by the gray bar. (C) Per-residue binding free energy difference calculated by subtracting the value of the residue in RBD_CoV_-bound ACE2 from that in RBD_CoV2_-bound ACE2. (D) Per-residue binding free energy difference calculated by subtracting the value of the residue in ACE2-bound RBD_CoV_ from that in ACE2-bound RBD_CoV2_.

On the side of RBDs, the residues with the binding free energy average values greater than 20 kJ/mol or lower than −20.0 kJ/mol include not only the charged residues but also the uncharged ones (Figure 4B). However, the absolute energy values of the charged residues (in a range of about 240-550 kJ/mol) are one order of magnitude greater than those of the uncharged residues (in a range of about 24-47 kJ/mol), with the positively and negatively charged ones making the positive and negative contributions to the binding affinities of both RBDs to ACE2, respectively, while the uncharged residues contributing positively to the ACE2 affinities of both RBDs. Interestingly, for the charged residues, those with the absolute values greater than 400 kJ/mol are all located at/near the binding interfaces (marked by black/dark gray rectangle), and those with absolute energy values lower than 300 kJ/mol are located distally from the binding interfaces (gray to lighter gray rectangles); while for the uncharged residues, they are all located at the RBD interface (marked by black rectangles).

Figure 4D shows the per-residue binding free energy difference from RBD_CoV_ to RBD_CoV2_. Clearly, all the significant differences occur on the residue substitutions involved in charge changes, with the loss of the negatively charged residues (i.e, E354N, D476G and D494S) and the gain of the positively charged residues (i.e., V417K, T444K, and H458K) resulting in the negative difference values and hence contributing to enhancing the ACE2 binding affinity of RBD_CoV2_ compared to that of RBD_CoV_, while the loss of the positively charged residues (i.e., R439N, K452L, K460N, and K478T) and the gain of the negatively charged residues (i.e., V471E and P484E) resulting in the positive difference values and contributing to reducing the ACE2 affinity of RBD_CoV2_ compared to that of RBD_CoV_. For all the above RBD residue substitutions, their significant differences in the binding free energy arise from the changes in the electrostatic interactions with ACE2 (Table S1), irrespective of the structural locations of substitutions. For example, E354N, despite being distal from the binding interfaces, leads to a significant loss in the electrostatic repulsive interactions with ACE2 (305.7 vs. 0.8 kJ/mol) and hence contributes to enhancing the ACE2 affinity.

In summary, our binding free energy calculations reveal that i) the higher binding affinity of RBD_CoV2_-ACE2 than of RBD_CoV_-ACE2 primarily originates from the significantly strengthened electrostatic attractive forces between RBD_CoV2_ and ACE2 than between RBD_CoV_ and ACE2, ii) the negatively charged residues in ACE2 and positively charged residues in RBDs make considerable positive contributions to the binding affinity due to their strong electrostatic attractive interactions with the other binding partners, while the positively charged residues in ACE2 and negatively charged residues in RBDs make considerable negative contributions to the binding affinity due to their strong electrostatic repulsive interactions with the other partners, iii) for the charged residues, their magnitudes of contributions to the binding affinity exhibit a trend of dependence on the residue’s distance to the binding interfaces; iv) the RBD residue substitutions involved in charge changes can greatly impact binding affinities of RBDs to ACE2 through altering the strength of electrostatic interactions with ACE2.

### 3.4 Interface residue contact networks and the related interactions

In order to further investigate how the RBD residue substitutions and complex structural fluctuations affect the interface interactions and further the protein-protein binding affinity, IRCNs were constructed over the representative structures of the two complexes, and the special interactions involved in IRCNs, including HBs and SBs, were also identified (Figure 5). The IRCN of RBD_CoV2_-ACE2 (Figure 5B) contains more nodes and edges than that of RBD_CoV_-ACE2 (Figure 5A), i.e., 39 (18 from RBD_CoV2_ and 21 from ACE2) and 37 (18 from RBD_CoV_ and 19 from ACE2) nodes, and 42 and 34 edges in the former and latter IRCNs, respectively, which result in the ratios of the number of edges to that of nodes being 1.08 and 0.92, respectively, indicating that a single residue forms on average more close interactions across the interfaces of RBD_CoV2_-ACE2 than of RBD_CoV_-ACE2. The above observations, together with a higher average number of inter-atomic close contacts between ACE2 and RBD_CoV2_ (142.3 ± 20.5) than between ACE2 and RBD_CoV_ (127.0 ± 19.0), reveal a tighter interface packing and more intensive interface interactions in RBD_CoV2_-ACE2 than in RBD_CoV_-ACE2 complex.

**Figure 5.**
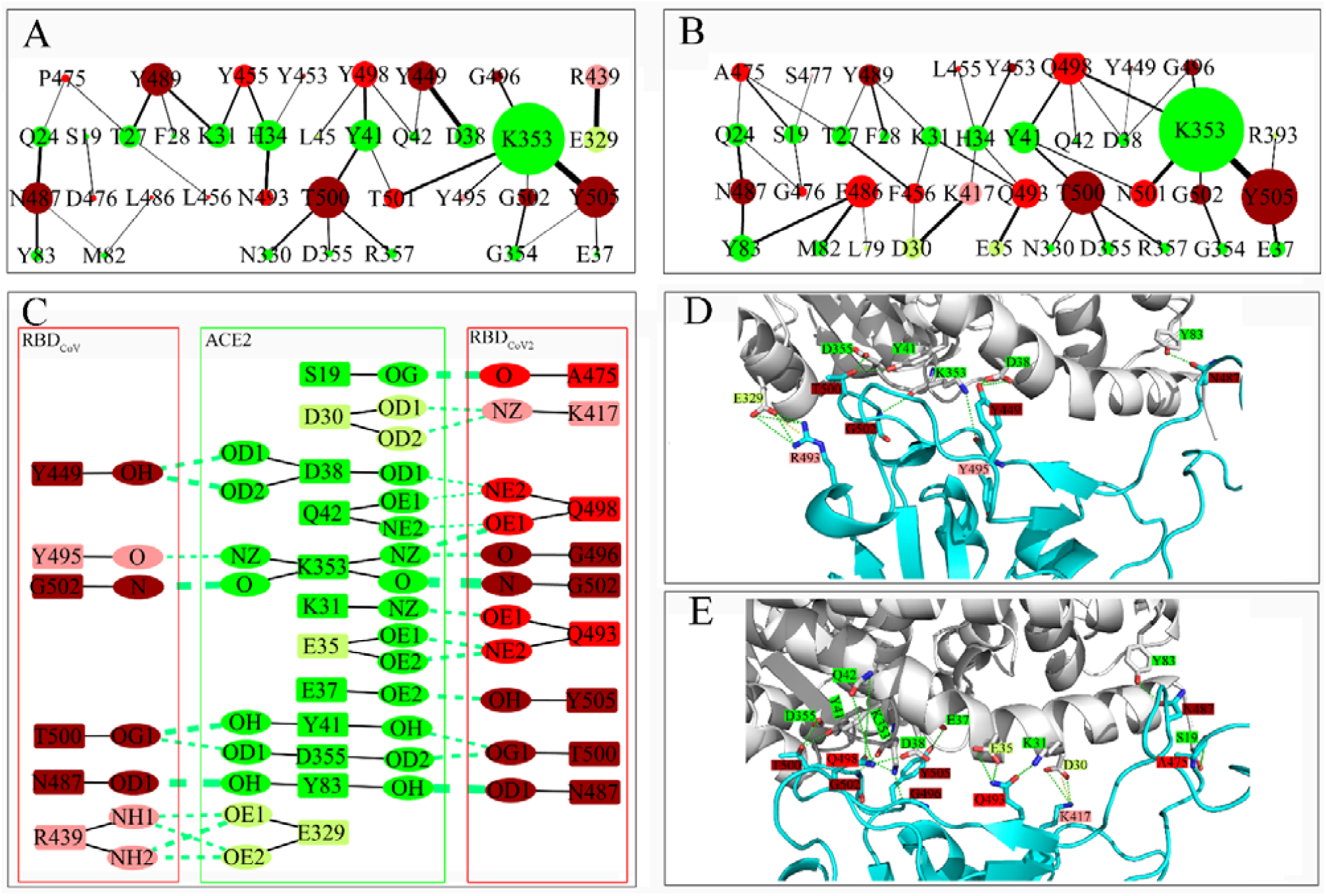
Interface residue contact networks (IRCNs) and special interactions across the binding interfaces of the RBD_CoV_-ACE2 and RBD_CoV2_-ACE2 complexes. (A) and (B) IRCNs constructed over the 100 representative structures of the RBD_CoV_-ACE2 and RBD_CoV2_-ACE2 complexes, respectively. Nodes are colored as follows: shared ACE2 residues between the two IRCNs: green; non-shared ACE2 residues: light green; conserved RBD residues between the two IRCNs: dark red; non-conserved/substituted RBD residue: red; non-shared RBD residues: light red. The node size represents the total number of close inter-atomic contacts occurring on a residue/node. The edge weight represents the average number of close inter-atomic contacts between the connected residues/nodes. (C) Interface hydrogen bonds (HBs) with occupancy greater than 20% identified over the respective 100 representative structures of the two complexes. HB-forming residues are marked using the same color scheme as in (A) and (B). HBs are represented by green dash lines connecting between the donor and acceptor atoms, with the line thickness representing the degree of the HB occupancy. (D) and (E) Structural locations of the interface HBs and salt bridges (SBs) in RBD_CoV_-ACE2 and RBD_CoV2_-ACE2, respectively. The representative structures of the two complexes are shown as cartoon representations, with ACE2 and RBDs colored gray and cyan, respectively. The residues involved in the formation of HBs and SBs are rendered as stick models, with oxygen and nitrogen atoms colored red and blue, respectively. HBs and SBs are represented by green and yellow dashed lines, respectively. Residue labels are colored the same as in (A) and (B).

There are eight conserved RBD residues (dark red nodes) between the two IRCNs (Figures 5A and B), out of which Y489, Y453, T500, and G502 have their respective identical local topologies due to the involvement of the same pairwise nodes, while the respective local topologies of Y449, G496, N487, and Y505 are slightly different between the two networks due to the gain/loss of close contacts with the surrounding ACE2 residues. Although the sizes of these individual conserved nodes can be larger or smaller between the two networks, the cumulative number of close contacts on them is slightly higher in the IRCN of RBD_CoV_-ACE2 (73.5) than in that of RBD_CoV2_-ACE2 (68.1); furthermore, the cumulative values of the vdW interaction energy, electrostatic interaction energy, and binding free energy are −72.8 and −68.9, −102.7 and −94.6, and −119.6 and −109.2 kJ/mol (Table S2), respectively, revealing a trend that the overall increased number of close contacts on the conserved RBD_CoV_ residues strengthen their vdW and electrostatic interactions with ACE2 and, hence, result in a larger positive contribution to the ACE2 binding affinity compared to the conserved RBD_CoV2_ residues.

With the exception of RBD:Y/L455, all the non-conserved RBD residues (red nodes) have larger node sizes in the IRCN of RBD_CoV2_-ACE2 than in that of RBD_CoV_-ACE2. The largest increase in the node size was observed for L486F (residue substitution from RBD_CoV_:L486 to RBD_CoV2_:F486; Figures 5A and B), which also leads to a more favorable contribution to the ACE2 binding affinity by −10.2 kJ/mol (Tables S1 and S2); since both RBD_CoV_:L486 and RBD_CoV2_:F486 are non-polar amino acids with their main chains forming no HB with the ACE2 residue, the enhanced ACE2 binding affinity by L486F mainly arises from the increased or additional vdW contacts/interactions with ACE2:M82, Y83, and L79 (Figures 5A and B, Table S2). The similar situation also occurs on L456F, which ranks the second in terms of the increased node size and enhances the ACE2 binding affinity by −4.9 kJ/mol. The third ranked increase in the node size occurs for N493Q, which considerably enhances the binding affinity by −36.7 kJ/mol; the reason for this is that the additional close contacts of RBD_CoV2_:Q493 with ACE2:K31 and E35 involve the formation of three stable HBs (Figures 5C and E), which strengthen the electrostatic interactions by −49.9 kJ/mol (Table S2). Y498Q also largely increases the node size and the ACE2 binding affinity (by −17.7 kJ/mol) because the formation of multiple additional HBs by RBD_CoV2_:Q498 with ACE2:D38, Q42, and K353 considerably strengthens the electrostatic interactions by −33.4 kJ/mol. P475A increases the node size due to the formation of a high-occupancy HB between RBD_CoV2_:A475 and ACE2:S19 (Figure 5C), which strengthens the electrostatic interactions by −17.3 kJ/mol. T501N and D476G lead to a limited increase in the node size. D476G leads to an abnormal enhancement in the ACE2 binding affinity by −337.9 kJ/mol (Figures 4D), with the reason being loss of the long-range electrostatic repulsive interactions rather than the slightly increased close contacts with ACE2 residues. Overall, the cumulative contacts of the non-conserved RBD interface residues with ACE2 are significantly more in the IRCN of RBD_CoV2_-ACE2 (63.7 and 60.2 with and without RBD_CoV2_:G476, respectively) than in that of RBD_CoV_-ACE2 (41.3 and 39.1 with and without RBD_CoV_:D476, respectively); in the RBD_CoV_-ACE2 complex, their cumulative binding free energy values with and without RBD_CoV_:D476 are 248.3 and −82.3 kJ/mol, respectively, and in the RBD_CoV2_-ACE2 complex the corresponding values with and without RBD_CoV2_:G476 are −165.9 and −158.6 kJ/mol, respectively, indicating that although D476G plays an overwhelming role in enhancing the ACE2 affinity, there is a significant trend that the increased close contacts on the non-conserved RBD_CoV2_ residues considerably enhance the ACE2 binding affinity of RBD_CoV2_, for which the strengthened electrostatic interactions arising from the formation of additional HBs by RBD_CoV2_:Q493 and Q498 with ACE2 play a dominant role.

There are non-shared nodes, either from RBDs (light red) or from ACE2 (light green), between the two IRCNs (Figures 5A and B). Two non-shared nodes, RBD_CoV_:R439 and RBD_CoV2_:K417, are worth noting. Specifically, the direct contacts of RBD_CoV_:R439 with ACE2:E329 are absent in the IRCN of RBD_CoV2_-ACE2 due to the R439N substitution, which results in losses of the direct HB and SB interactions and the indirect long-range electrostatic forces of attraction present in RBD_CoV_-ACE2 (Figures 5C and D) and, hence, reduces the ACE2 binding affinity by 547.0 kJ/mol (Table S2). The residues at position 417 are RBD_CoV2_:K417 and RBD_CoV_:V417. Although both residues are located outside the RBM region, RBD_CoV2_:K417 forms close contacts with ACE2:D30 and H34 due to its longer, positively charged side chain compared to that of RBD_CoV_:V417. In particular, the close contacts of RBD_CoV2_:K417 with ACE2:D30 involve the formation of two HBs and one SB (Figures 5C and E), which together with the long-range electrostatic forces of attraction to ACE2, provide the most favorable contribution to the ACE2 binding affinity (−495.1 kJ/mol) among all the RBD_CoV2_ residues (Figure 4B and Table S1); also worth noting is that, among all residue substitutions, V417K most significantly enhances the ACE2 binding affinity of RBD_CoV2_ (by −487.6 kJ/mol; Figure 4D). The other two non-shared RBD nodes, RBD_CoV_:Y495 and RBD_CoV2_:S477, contribute limitedly to enhancing the ACE2 affinities of respective RBDs through slightly strengthening their vdW and electrostatic interactions with ACE2 compared to the structurally equivalent residues. Overall, the cumulative binding free energy values of the non-shared RBD residues in IRCNs of RBD_CoV_-ACE2 and RBD_CoV2_-ACE2 are −560.6 and −504.1 kJ/mol (Table S2), respectively, indicating that the non-shared RBD residues are more conducive to enhancing the ACE2 binding affinity of RBD_CoV_; nevertheless, when taking all the RBD interface residues (including the conserved, non-conserved, and non-shared RBD nodes) into account, their cumulative binding free energy values are −431.9 and −779.2 kJ/mol, respectively (Table S2), indicating that the interface residues of RBD_CoV2_ are more conducive to enhancing the ACE2 binding affinity than those of RBD_CoV_.

On the side of ACE2, the non-shared nodes L45 and E329 in the IRCN of RBD_CoV_-ACE2 enhance the binding affinity of ACE2 to RBD_CoV_ compared to that to RBD_CoV2_ by −4.7 and −75.4 kJ/mol (Table S3), respectively. As a matter of fact, ACE2:E329 simultaneously contributes positively to ACE2’s affinity to both RBD_CoV_ (−158.7 kJ/mol) and RBD_CoV2_ (−83.2 kJ/mol) due to its electronegativity; nevertheless, its larger positive contribution to the RBD_CoV_ binding affinity can be attributed to the formation of close contacts, in particular of the four stable HBs and one SB with RBD_CoV_:R439 (Figures 5C and D), which are absent in the IRCN of RBD_CoV2_-ACE2 due to the R439N substitution. In the IRCN of RBD_CoV2_-ACE2 there are four non-shared ACE2 nodes, i.e., L79, D30, E35, and R393, out of which L79 is a non-polar residue and hence, its close contacts with RBD_CoV2_:F486 enhances, to a limited degree, the ACE2’s binding affinity to RBD_CoV2_ (−5.2 kJ/mol) compared to that to RBD_CoV_ (−1.8 kJ/mol), while the rest three residues are charged ones that contribute considerably differently to the binding affinity of ACE2 to the two RBDs, depending on their charge properties and/or the residue type of the RBD node to which they are connected. Specifically, the negatively charged ACE2:D30 makes only few contacts (lightly weighted edge) with the hydrophobic RBD_CoV2_:F456 while quite a few contacts (heavily weighted edge) with the positively charged RBD_CoV2_:K417 (Figure 5B) that involve the formation of HBs and SB as described above, which significantly strengthen the electrostatic attractive interactions of ACE2:D30 with RBD_CoV2_ (−226.5 kJ/mol) compared to those with RBD_CoV_ (−66.5 kJ/mol), thus greatly enhancing the binding affinity to RBD_CoV2_ (−155.7 kJ/mol) compared to that to RBD_CoV_ (−64.1 kJ/mol; see Table S3). On the contrary, although the negatively charged ACE2:E35 is connected to RBD_CoV2_:Q493 by a heavily weighted edge and also forms HBs with RBD_CoV2_:Q493, ACE2:E35 only moderately enhances the affinity to RBD_CoV2_ (−59.0 kJ/mol) compared to that to RBD_CoV_ (−46.5 kJ/mol). The reason for this is that RBD_CoV2_:Q493 is a polar uncharged residue without carrying a net positive charge and hence, cannot form SB with ACE2:E35. ACE2:R393 makes considerable negative contributions to the binding affinity of ACE2 to both RBD_CoV_ and RBD_CoV2_ due to its electropositivity, with the calculated binding free energy values being of 72.4 and 100.4 kJ/mol, respectively (Table S3). However, only in the IRCN of RBD_CoV2_-ACE2 can ACE2:R393 make close contacts with RBD_CoV2_:Y505, which although result in a slightly more negative vdW interaction energy (−1.9 kJ/mol) than in RBD_CoV_-ACE2 (−1.1 kJ/mol), bring about much more positive electrostatic interaction energy (110.3 kJ/mol) than in RBD_CoV_-ACE2 (75.3 kJ/mol; see Table S3), indicating that the larger negative contribution of ACE2:R393 to the binding affinity to RBD_CoV2_ arises from the much stronger electrostatic forces of repulsion due to its closer distance to RBD_CoV2_. Despite more non-shared ACE2 nodes in the IRCN of RBD_CoV2_-ACE2, their cumulative binding free energy value (−119.6 kJ/mol) is less negative than that of the non-shared ACE2 nodes in the IRCN of RBD_CoV_-ACE2 (−164.4 kJ/mol); however, when taking the shared ACE2 residues into account, the cumulative binding free energy values reach −308.5 and −246.8 kJ/mol, respectively, indicating that the ACE2 interface residues make an overall larger positive contribution to the binding affinity to RBD_CoV2_ than to RBD_CoV_.

In summary, the RBD_CoV2_-ACE2 complex has more inter-atomic close contacts across the binding interfaces and a tighter interface packing than RBD_CoV_-ACE2. Although the overall increased number of interface close contacts strengthen the favorable vdW interactions between RBD_CoV2_ and ACE2 (Tables S2 and S3), the enhanced binding affinity of RBD_CoV2_-ACE2 is primarily determined by the significantly strengthened electrostatic attractive interactions occurring on its interface residues, in which the gain of HBs and SB as well as of the long-range electrostatic forces of attraction due to the V417K substitution, the loss of the electrostatic forces of repulsion due to the D476G substitution, the gain of HBs due to the substitutions of N493Q and Y498Q, and the increased number of HBs formed by the ACE2 interface residues with RBD_CoV2_, play crucial roles.

## 4. Discussion

Although the experimentally determined crystal structures of RBD_CoV_-ACE2 and RBD_CoV2_-ACE2 [13,14] show overall similar conformations and nearly identical protein-protein binding modes, our MD simulation-based analyses reveal distinctly different dynamic and thermodynamic behaviors between them. When compared to the RBD_CoV2_-ACE2 complex, the RBD_CoV_-ACE2 complex features the enhanced global structural fluctuations and has larger conformational entropy and conformational diversity. Of note is that during MD simulations the individual bound proteins also underwent larger structural fluctuations in RBD_CoV_-ACE2 than in RBD_CoV2_-ACE2, and in particular, the two binding partners experienced larger relative movements with respect to each other in the former complex, suggesting a correlation that the enhanced protein-protein association could enhance the structural stability of both the individual binding partners and the entire complex. Undoubtedly, it is the amino-acid-sequence differences between RBD_CoV_ and RBD_CoV2_ and the resultant changes in RBD physicochemical properties and RBD-ACE2 interaction strengths that give rise to the observed different dynamic and thermodynamic behaviors of the two complexes, which in turn may also have impacts on interactions and binding affinities between RBDs and ACE2.

The detailed comparison of the calculated values of various MM-PBSA energy terms (Table 1) reveals that i) the inter-protein electrostatic interaction strength/energy determines the high-affinity bindings of both RBDs to ACE2 and, ii) the significantly strengthened electrostatic attractive interactions between RBD_CoV2_ and ACE2 determine the higher binding affinity of RBD_CoV2_-ACE2 than of RBD_CoV_-ACE2. ACE2 is heavily negatively charged at pH 7.4 with the predicted net charges of −25 and −24 in the RBD_CoV_-ACE2 and RBD_CoV2_-ACE2 complexes, respectively, thus generating an overall very strong electronegative potential around itself (Figure S3); whereas both RBD_CoV_ and RBD_CoV2_ have a net charge of +2, thus generating an overall moderate electropositive potential while with different distributions of the local electropositive and electronegative potentials (Figure S3). Consequently, ACE2 has strong electrostatic forces of attraction to both RBDs, while the observed difference in the inter-protein electrostatic interaction strengths of the two complexes could be attributed to the different distributions of the electropositive and electronegative potentials on the two RBDs, which in turn depend on the structural locations of the positively and negatively charged residues. Therefore, it is reasonable to speculate a scenario in which the strong electronegative potential around ACE2 could accelerate the diffusion of both net positively charged RBDs toward ACE2 through a strong electrostatic attraction and hence facilitate the initial recognition and subsequent orientation adjustment between them, while different electrostatic interaction strengths resulting from different charged residue distributions on RBD_CoV_ and RBD_CoV2_ determine the final difference in binding affinities of the two RBDs to ACE2.

A previous study [57] on the electrostatic features of the two complexes reveals that, although both RBDs are attractive to ACE2 by strong electrostatic forces, the RBD residue substitutions involved in charge changes and the resulting differences in the distributions of the charged residues and hence the related electric field line density determine the stronger electrostatic attractive forces of ACE2 to RBD_CoV2_ than to RBD_CoV_, thus supporting our speculation. In the current study, we found that, although the charged residues commonly exhibit large binding free energy values, there is a trend that the magnitudes of the absolute energy values largely depend on the distance of the charged residues to the binding interfaces (Figures 4A and B). In fact, it is the distance-dependent changes in the electrostatic interaction strength of the charged residues that give rise to their different magnitudes of contribution to the binding affinity (Figure S4). Although with exceptions, a general trend is observable that the closer the distance of the negatively charged ACE2 residues and positively charged RBD residues to the binding interfaces, the stronger the electrostatic attractive interactions with the other partners (Figures S4C, D, and F), and hence the greater the positive contribution to the affinity (Figures S4A, B, and E); a similar trend occurs for the positively charged ACE2 residues and negatively charged RBD residues in which the electrostatic repulsive interactions increase as the distance to the interfaces decreases.

Comparison between the constructed IRCNs of the two complexes reveals that RBD residue substitutions lead to more intensive interface contacts and a tighter interface packing in the RBD_CoV2_-ACE2 than in the RBD_CoV_-ACE2 complex, which on the one hand explain the reduced inter-protein positional movements in RBD_CoV2_-ACE2, and on the other hand, significantly enhance the overall electrostatic interaction strength of the interface residues with the other partner in RBD_CoV2_-ACE2. For RBD_CoV2_-ACE2 and RBD_CoV_-ACE2, the values of the inter-protein electrostatic interaction energy are −2597.0 and −2338.4 kJ/mol (Table 1), respectively, the cumulative values of the electrostatic interaction energy of all the IRCN-forming residues (i.e., the interface residues) are −1258.6 and −799.8 kJ/mol (Tables S2 and S3), respectively, and those of all the non-interface residues are −1338.4 and −1538.7 kJ/mol, respectively. Therefore, despite the importance of the long-range indirect electrostatic attractive interactions in promoting the high-affinity bindings of both RBDs to ACE2, the stronger electrostatic attractive forces occurring on the interface residues of RBD_CoV2_-ACE2 dictate the overall stronger inter-protein electrostatic interactions of RBD_CoV2_-ACE2 than of RBD_CoV_-ACE2. Further comprehensive comparative analyses of IRCNs, IRCN-related interactions, and energy components of IRCN-forming residues between the two complexes reveal that the difference in the electrostatic interaction strength depends on the charge properties of the interface residues, the number of close contacts on the charged residues, and whether or not the close contacts involve the formation of HB and SB.

Of interest is that for the RBM residues (438-506) of the RBD_CoV2_ and RBD_CoV_, the cumulative values of the binding free energy are −574.3 and −1313.7 kJ/mol, respectively, and the cumulative values of electrostatic interaction energy are −538.7 and −1300.8 kJ/mol (Table S4), respectively; therefore, the stronger electrostatic interactions of RBD_CoV_ RBM with ACE2 determine its larger positive contribution to the ACE2 binding affinity than RBD_CoV2_ RBM. The net charges of RBMs in RBD_CoV_ and RBD_CoV2_ are +3 and +1, respectively, which explains why RBM has stronger electrostatic forces of attraction to ACE2 in RBD_CoV_-ACE2. Since i) the RBD interface residues (i.e., IRCN-forming residues from RBDs) identified in our work include only a fraction of RBM residues (Figure 1D), ii) the net charges of the RBD_CoV_ interface and RBD_CoV2_ interface are 0 (determined by R439 and D476; see Figure 5A) and +1 (determined by K417 that does not belong to RBM), respectively, and iii) RBD_CoV2_ interface residues form more HBs with ACE2 than RBD_CoV_ interface residues (Figure 5C), it is not surprising to observe that the RBD_CoV2_ interface has stronger electrostatic forces of attraction to ACE2 than the RBD_CoV_ interface. To this end, we could conclude that it is the effective RBD interface rather than RBM that dominates the higher ACE2 binding affinity of RBD_CoV2_ than of RBD_CoV_, suggesting that when estimating the changes in the ACE2 binding affinity between different RBDs or RBD mutants, one may first pay attention to the changes in the electrostatic interactions caused by the residue substitutions at the RBD interface, then those near the interface, and finally those located distally from the interface, rather than simply focusing on the residue substitutions in RBM.

Although we do not calculate the binding free energies between human ACE2 and RBDs of various variants of concern (VOC), our findings can still facilitate the explanation of the experimentally observed changes in the ACE2 binding affinities of different VOC RBDs from the perspective of the electrostatic interaction change principle. For example, RBD of the Delta variant (B.1.617.2 lineage) contains two residue substitutions, L452R and T478K [58], which result in the gain of two positively charged residues near the binding interfaces and hence could greatly strengthen the electrostatic forces of attraction to ACE2. Of interest is that in RBD_CoV_ the corresponding residues are positively charged K452 and K478, which compared to RBD_CoV2_:L452 and T478 enhance the ACE2 binding affinity of RBD_CoV_ (by −408.8 and −306.3 kJ/mol, respectively; see Figure 4D) through significantly strengthening their electrostatic forces of attraction to ACE2 (by −413.3 and −301.9 kJ/mol, respectively; see Table S4). These observations may help explain the experimentally observed 1.2-4.6-fold increase in the ACE2 affinity of the Delta RBD compared to the wild-type (WT) RBD_CoV2_ [59-61].

The recently emerged VOC Omicron (B.1.1.529 lineage) contains 37 residue substitutions in the spike protein, of which 15 are in RBD and 11 near/at the binding interfaces [62,63]. Of the four residue substitutions (G339D, S371L, S373P, S375F) with location far from the binding interfaces, G339D would largely reduce the ACE2 affinity due to the long-range electrostatic repulsion of D339 with ACE2 (as observed for E340 in both RBD_CoV_ and RBD_CoV2_; see Figure 4B), while the effects of the other three substitutions could be ignored. Of the 11 substitutions (K417N, N440K, G446S, S477N, T478K, E484A, Q493R, G496S, Q498R, N501Y, and Y505H) located near/at the RBD interface, six (K417N, N440K, T478K, E484A, Q493R, and Q498R) involve charge changes and could greatly impact the ACE2 affinity. Specifically, E484A would overcompensate for the negative effect of G339D because the distance of residue 484 to the binding interfaces is closer than that of residue 339 (Figure 4B) and, hence, E484A could lead to more loss of the electrostatic repulsion with ACE2 compared to the added electrostatic repulsion by G339D; although all the other five substitutions involve either the gain or loss of the positively charged residues, they lead to a net gain of three positive charges, thus increasing the number of net positive charges from +2 in WT RBD_CoV2_ to +5 in Omicron RBD. This, in conjunction with the close distances/contacts of all the newly acquired positively charged residues to/with ACE2, could greatly enhance the ACE2 binding affinity of the Omicron RBD compared to that of WT RBD_CoV2_. The above inference is confirmed by the experimental measurements showing that the Omicron RBD enhances the ACE2 affinity by 1.4-2.4 folds compared to WT RBD_CoV2_ [59-61]. In addition, a computational study [64] shows that the substitutions involving the gain of the positively charged residues (N440K, T478K, Q493R, and Q498R) and loss of the negatively charged residue (E484A) collectively enhance, although to different extents, the ACE2 binding affinity, and the gain of the negatively charged residue (G339D) reduces the ACE2 affinity to an extent that can be overcompensated for by the increased affinity contributed by E484A, in line with our electrostatic interaction change principle-based inference. Also worth noting is that a Cryo-EM study reveals [60] that Q493R and Q498R lead to the formation of new HBs and SBs with ACE2:E35 and D38, respectively, which can over offset the local electrostatic repulsion with ACE2:K31 and K353, respectively, ultimately resulting in a large increase in the electrostatic interaction strength with ACE2 and enhancing the affinity.

## 5. Conclusions

Through comprehensive comparative analysis of our computational results, we conclude that it is the RBD residue substitutions at the binding interface that lead to the overall stronger electrostatic force of attraction of ACE2 to RBD_CoV2_ than to RBD_CoV_, which on the one hand tightens the interface packing and reduces the structural fluctuations of RBD_CoV2_-ACE2, and on the other hand, enhances the binding affinity of RBD_CoV2_-ACE2 compared to that of RBD_CoV_-ACE2. We also highlight that i) although the RBD residue substitutions involved in charge changes can significantly impact the inter-protein electrostatic interaction strength and hence the binding affinity, the extent of the impact largely depends on the distance of the substitution site to the binding interfaces (i.e., the extent of the impact increases/decreases as the distance decreases/increases) and, ii) the formation or destruction of the interface HBs and SBs caused by RBD residue substitutions or interface conformational changes can also largely impact the inter-protein electrostatic interactions. Our findings not only shed light on the mechanical and energetic mechanisms responsible for modulating the inter-protein interaction strengths and binding affinities of the two RBD-ACE2 complexes, but can also help predict the binding affinity changes of different VOC RBDs with ACE2 and, further, help estimate the transmissibility/infectivity of newly emerged variants of SARS-CoV-2.

## Supplementary Materials

The following supporting information can be downloaded at: www.xxx.com, **Figure S1**: Time dependent C_α_ RMSD values of RBD and ACE2 after least-squares fitting to either RBD or ACE2 in the starting complex structure during the multiple-replica MD simulations; **Figure S2**: Eigenvalues of the first 30 eigenvectors (main plot) and cumulative contribution of all eigenvectors to the total mean square fluctuations (inset plot) for the RBD_CoV_-ACE2 (black line) and RBD_CoV-2_-ACE2 complexes (red line); **Figure S3**: 3D structures and electrostatic potential maps of the RBD_CoV_-ACE2 and RBD_CoV2_-ACE2 complexes; **Figure S4**: The trends of the binding free energy and electrostatic energy of the charged residues in RBD_CoV_-ACE2 and RBD_CoV2_-ACE2 at different distances to the binding interfaces; **Table S1**: Average values of the per-residue binding free energy and energy components in the RBD_CoV_-ACE2 and RBD_CoV2_-ACE2 complexes calculated over their respective 100 representative structures (see the file Table_S1.xlsx); **Table S2**: Values of the residue binding free energy and energy components (kJ/mol) for IRCN-forming residues from RBD_CoV_ and RBD_CoV2_; **Table S3**: Values of the residue binding free energy and energy components (kJ/mol) for the ACE2 residues that participate in the formation of IRCNs of the RBD_CoV_-ACE2 and RBD_CoV2_-ACE2 complexes; Table S4: Values of the residue binding free energy and energy components (kJ/mol) for RBM residues from RBD_CoV_ and RBD_CoV2_.

## Author Contributions

Conceptualization, Y.-X.F. and S.-Q.L.; methodology, Z.-B.Z. and Y-.L.X.; validation, Z.-B.Z., Y.-L.X., J-.X.S., and W.-W.D.; formal analysis, Z.-B.Z. and Y.-L.X.; investigation, Z.-B.Z., Y.-L.X., J-.X.S., and W.-W.D.; data curation, Z.-B.Z. and Y-.L.X.; writing—original draft preparation, Z.-B.Z.; writing—review and editing, S.-Q.L. and Y.-X.F.; visualization, Z.-B.Z. and Y.-L.X.; supervision, Y.-X.F. and S.-Q.L.; project administration, S.-Q.L.; funding acquisition, Y.-X.F. and S.-Q.L.. All authors have read and agreed to the published version of the manuscript.

## Institutional Review Board Statement

Not applicable.

## Informed Consent Statement

Not applicable.

## Data Availability Statement

All data are contained within the article or its supplementary materials as Figures or Tables.

