## Supplementary for "Mechanistic origin of different binding affinities of SARS-CoV and SARS-CoV-2 spike RBDs to human ACE2"

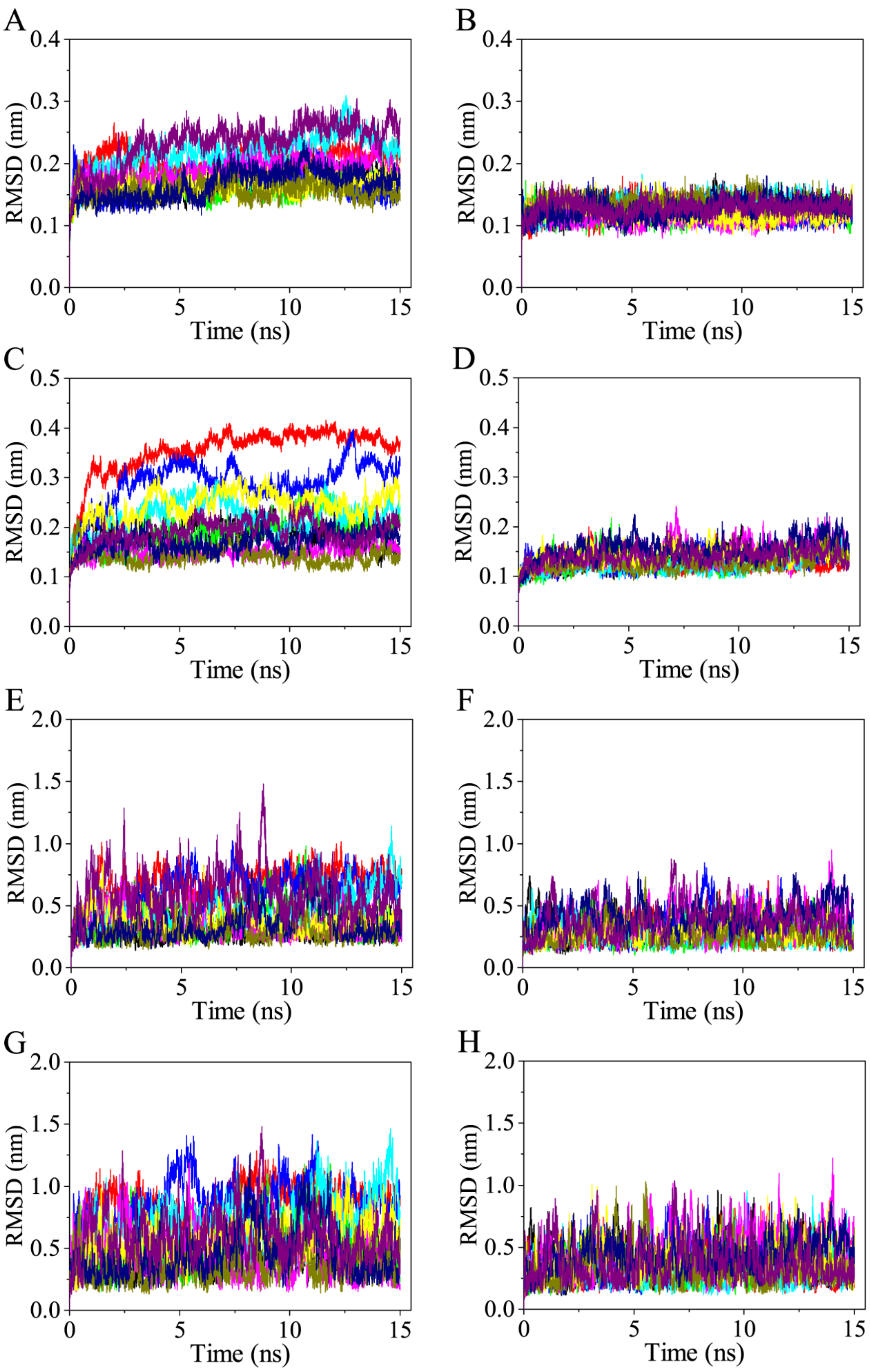


**Figure S1.** Time dependent $C_\alpha$ RMSD values of RBD and ACE2 after least-squares fitting to either RBD or ACE2 in the starting complex structure during the multiple-replica MD simulations. (A) and (B) RMSD curves of $RBD_{CoV}$ and $RBD_{CoV2}$ after least-squares fitting to the respective RBD structures in the starting complexes, respectively. (C) and (D) RMSD curves of ACE2 after least-squares fitting to the respective ACE2 structures in the starting complexes of $RBD_{CoV}$-ACE2 and $RBD_{CoV2}$-ACE2, respectively. (E) and (F) RMSD curves of $RBD_{CoV}$ and $RBD_{CoV2}$ after least-squares fitting to the ACE2 structure in the respective starting complexes, respectively. (G) and (H) RMSD curves of ACE2 after least-squares fitting to the structures of $RBD_{CoV}$ and $RBD_{CoV2}$ in the respective starting complex, respectively.

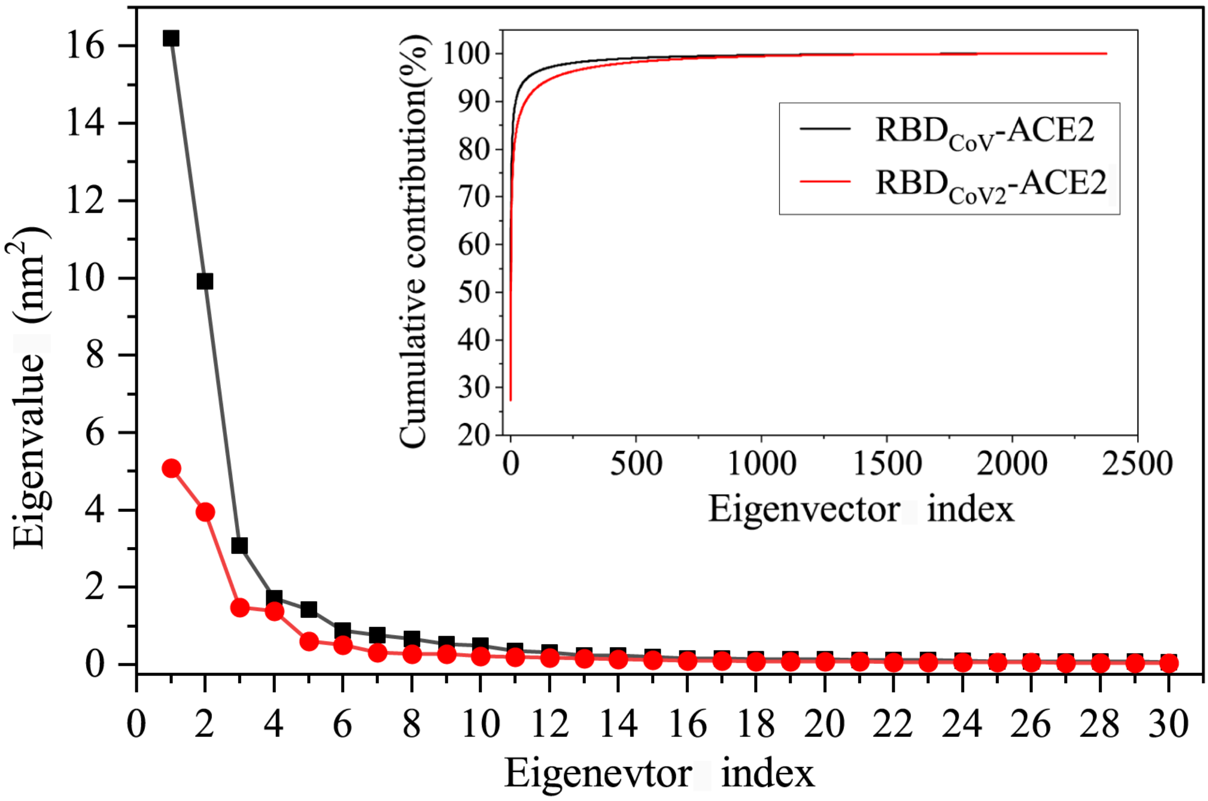


**Figure S2.** Eigenvalues of the first 30 eigenvectors (main plot) and cumulative contribution of all eigenvectors to the total mean square fluctuations (inset plot) for the RBD$_{CoV}$-ACE2 (black line) and RBD$_{CoV-2}$-ACE2 complexes (red line).

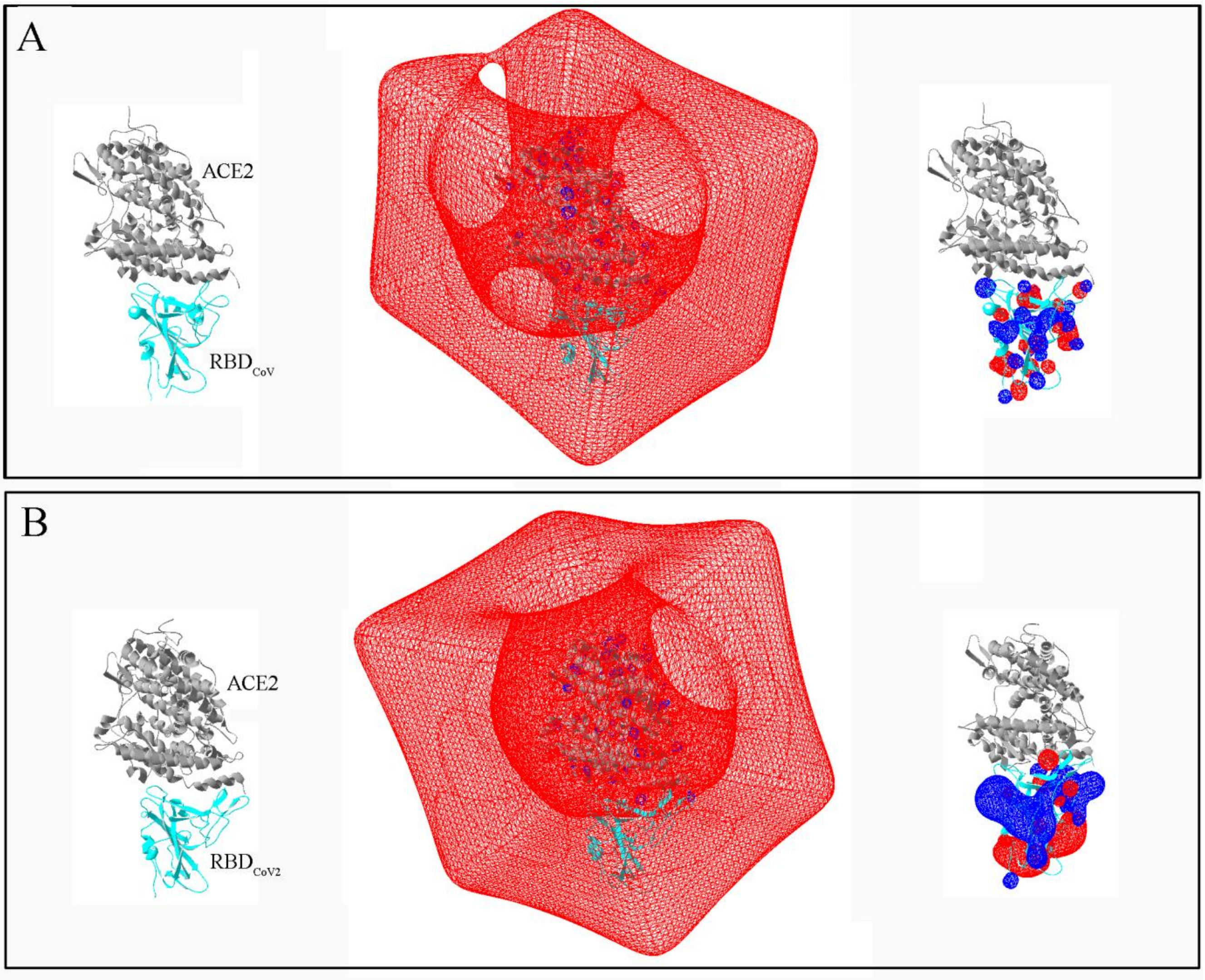


**Figure S3.** 3D structures and electrostatic potential maps of the $RBD_{CoV}$-ACE2 and $RBD_{CoV2}$-ACE2 complexes. (A) $RBD_{CoV}$-ACE2. (B) $RBD_{CoV2}$-ACE2. In (A) and (B), the left, middle, and right panels show the cartoon representation of a randomly selected representative complex structure, electrostatic potential map of ACE2, and electrostatic map of RBD, respectively. In the middle and right panels, the red and blue contours denote the electronegative potential (–10) and electropositive potential (+10), respectively. It is clear that ACE2s in both complexes have a very strong electronegative potential that extends beyond the ACE2 surface and wrap RBDs completely, whereas the two RBDs feature distinctly different distributions of the electropositive and electronegative potentials, i.e., mixed distributions of small local electropositive and electronegative potentials over $RBD_{CoV}$, while the relatively concentrated distributions of the large clumpy electropositive and electronegative potentials on the middle upper part and lower part of $RBD_{CoV2}$, respectively, although several small local electronegative potentials are located near the binding interfaces. The electrostatic potential maps were generated by Swiss-PdbViewer [1].

1. Guex, N.; Peitsch, M.C. SWISS-MODEL and the Swiss-PdbViewer: an environment for comparative protein modeling. Electrophoresis 1997, 18, 2714-2723, doi:10.1002/elps.1150181505.

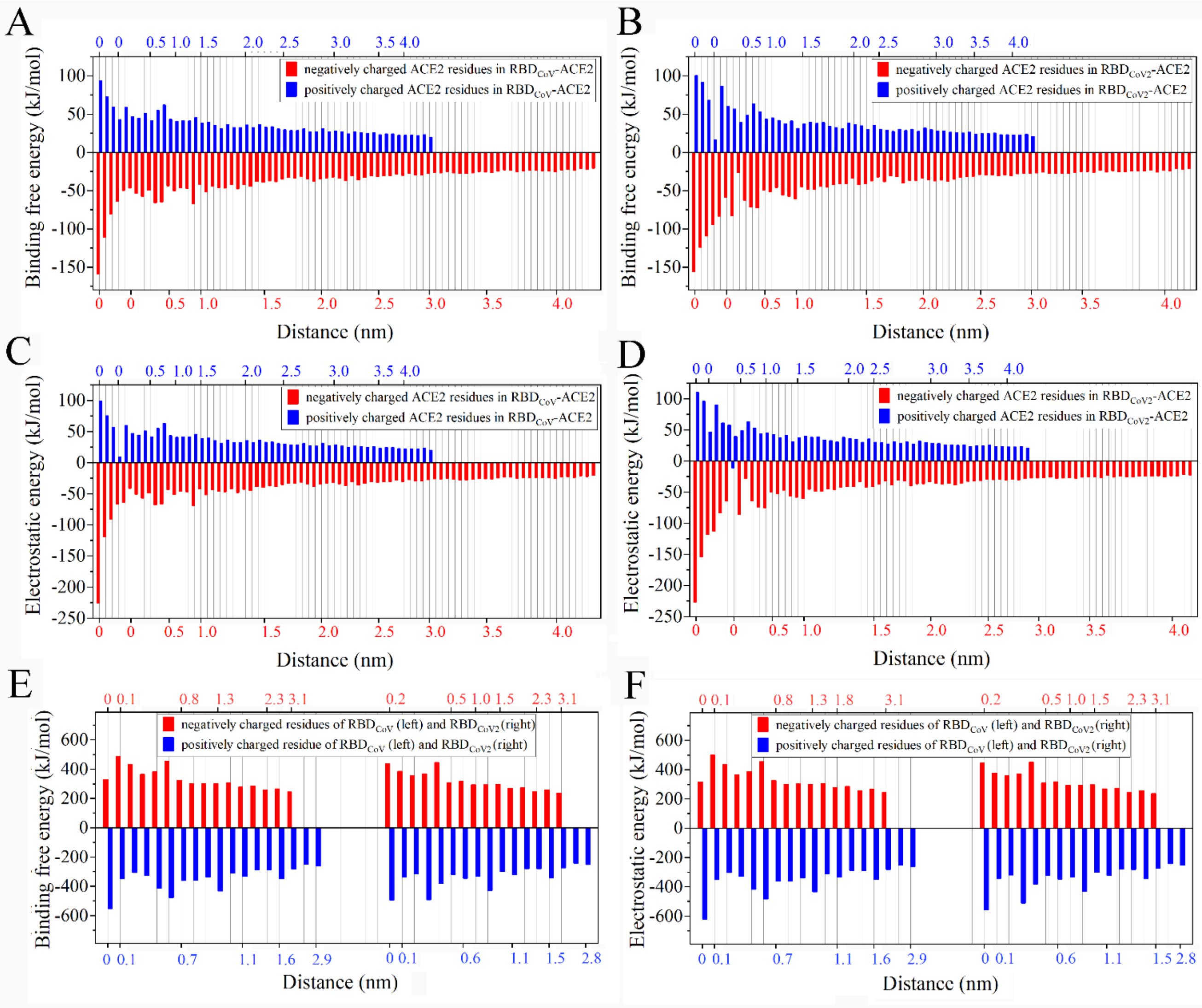


**Figure S4.** The trends of the binding free energy and electrostatic energy of the charged residues in $RBD_{CoV}$-ACE2 and $RBD_{CoV2}$-ACE2 at different distances to the binding interfaces. (A) and (B) Binding free energy average values of the positively charged (blue) and negatively charged (red) ACE2 residues in the $RBD_{CoV}$-ACE2 and $RBD_{CoV2}$-ACE2 complexes, respectively. (C) and (D) Average values of the electrostatic interaction energy of the charged ACE2 residues with $RBD_{CoV}$ and $RBD_{CoV2}$, respectively. (E) Binding free energy average values of the charged residues from $RBD_{CoV}$ (left) and $RBD_{CoV2}$ (right). (F) Average values of the electrostatic interaction energy of the charged residues of $RBD_{CoV}$ (left) and $RBD_{CoV2}$ (right) with ACE2. The charged residues with distance of 0 nm are the interface residues identified in this work. Note that there are more than one charged interface residues in the two complexes, and therefore their absolute energy values are sorted in descending order while with the distances labeled as 0. The actual distances of the non-interface charged residues to the binding interfaces are labeled along either the top or bottom horizontal axis.

**Table S1.** Average values of the per-residue binding free energy and energy components in the $RBD_{CoV}$-ACE2 and $RBD_{CoV2}$-ACE2 complexes calculated over their respective 100 representative structures (see the file **Table_S1.xlsx**).

**Table S2**. Values of the residue binding free energy and energy components (kJ/mol) for IRCN-forming residues from $RBD_{CoV}$ and $RBD_{CoV2}$.

| $RBD_{CoV}$ | | | | | | $RBD_{CoV2}$ | | | | | |
|---|---|---|---|---|---|---|---|---|---|---|---|
| Residue[a] | $\triangle E_{vdW}$ | $\triangle E_{elec}$ | $\triangle G_{polar}$ | $\triangle G_{non\text{-}polar}$ | $\triangle G_{binding}$ | Residue[a] | $\triangle E_{vdW}$ | $\triangle E_{elec}$ | $\triangle G_{polar}$ | $\triangle G_{non\text{-}polar}$ | $\triangle G_{binding}$ |
| V417 | –0.8 | -6.7 | 0.1 | -0.1 | -7.5 | K417 | -1.1 | -557.3 | 64.2 | -0.9 | -495.1 |
| R439 | 1.5 | -619.8 | 67.9 | -1.0 | -551.4 | N439 | -0.4 | -3.6 | -0.4 | 0.0 | -4.4 |
| Y449 | -3.1 | -34.2 | 10.5 | -1.1 | -27.8 | Y449 | -5.1 | 1.9 | 3.0 | -0.8 | -1.0 |
| Y453 | -3.2 | 2.7 | 1.9 | -0.4 | 1.0 | Y453 | -3.4 | -1.9 | 4.6 | -0.5 | -1.2 |
| Y455 | -12.0 | 3.4 | 5.6 | -1.5 | -4.6 | L455 | -10.6 | 7.8 | -1.7 | -0.9 | -5.5 |
| L456 | -4.3 | -2.1 | 0.2 | -0.5 | -6.8 | F456 | -10.1 | -2.8 | 2.4 | -1.2 | -11.7 |
| P475 | -11.4 | -2.4 | 2.9 | -1.6 | -12.5 | A475 | -5.2 | -19.7 | 11.6 | -1.2 | -14.5 |
| D476 | -3.0 | 318.0 | 16.3 | -0.7 | 330.6 | G476 | -4.3 | -4.7 | 2.1 | -0.4 | -7.3 |
| G477 | -0.3 | -4.2 | 0.1 | 0.0 | -4.3 | S477 | -2.5 | -9.5 | 3.5 | -0.5 | -9.0 |
| L486 | -5.4 | -8.6 | 1.2 | -1.2 | -14.0 | F486 | -15.6 | -11.7 | 5.3 | -2.2 | -24.2 |
| N487 | -9.3 | -15.5 | 14.8 | -1.7 | -11.7 | N487 | -7.6 | -20.6 | 14.1 | -0.9 | -15.1 |
| Y489 | -16.8 | -5.0 | 7.5 | -2.0 | -16.2 | Y489 | -14.7 | 0.8 | 8.0 | -1.8 | -7.7 |
| N493 | -7.1 | -11.3 | 9.5 | -1.0 | -9.9 | Q493 | -7.5 | -61.2 | 23.9 | -1.7 | -46.6 |
| Y495 | -2.1 | -13.2 | 6.4 | -0.3 | -9.2 | Y495 | -1.8 | -7.3 | 0.8 | 0.0 | -8.3 |
| G496 | -3.0 | -10.0 | 5.5 | -0.2 | -7.7 | G496 | -1.7 | -19.9 | 6.6 | -0.6 | -15.6 |
| Y498 | -14.6 | -0.7 | 4.2 | -1.6 | -12.6 | Q498 | -8.1 | -34.1 | 13.2 | -1.3 | -30.3 |
| T500 | -13.8 | -17.8 | 14.5 | -2.1 | -19.3 | T500 | -12.9 | -13.1 | 11.7 | -2.3 | -16.6 |
| T501 | -14.0 | -12.5 | 5.3 | -0.9 | -22.1 | N501 | -15.2 | -24.4 | 14.5 | -0.8 | -25.9 |
| G502 | -6.5 | -9.1 | 4.0 | -1.0 | -12.6 | G502 | -5.1 | -10.0 | 3.5 | -0.8 | -12.3 |
| Y505 | -17.1 | -13.8 | 7.3 | -1.7 | -25.3 | Y505 | -18.5 | -31.7 | 12.9 | -2.3 | -39.6 |
| **Sum1** | -72.8 | -102.7 | 66.2 | -10.3 | -119.6 | **Sum1** | -68.9 | -94.6 | 64.3 | -10.0 | -109.2 |
| **Sum2** | -71.7 | 283.8 | 16.3 | -0.7 | 248.3 | **Sum2** | -76.7 | -150.8 | 71.1 | -9.6 | -165.9 |
| **Sum3** | -0.6 | -633.0 | 74.3 | -1.3 | -560.6 | **Sum3** | -3.6 | -566.8 | 67.7 | -1.4 | -504.1 |
| **Sum4** | -145.1 | -451.9 | 185.7 | -20.7 | -431.9 | **Sum4** | -149.1 | -812.2 | 203.1 | -20.9 | -779.2 |

[a] RBD interface residues (i.e., IRCN-forming residues from RBDs) are colored the same as in Figures 5A and B in the main body of the paper, with the conserved, non-conserved, and non-shared residues colored dark red, red, and light red, respectively; the non-interface RBD residues (i.e., those do not make close contact with ACE2) are colored black; Sum1-4 indicate the cumulative values of respective energy terms for the conserved, non-conserved, non-shared, and all RBD interface residues, respectively.

**Table S3**. Values of the residue binding free energy and energy components (kJ/mol) for the ACE2 residues that participate in the formation of IRCNs of the $RBD_{CoV}$-ACE2 and $RBD_{CoV2}$-ACE2 complexes.

| $RBD_{CoV}$-bound ACE2 | | | | | | $RBD_{CoV2}$-bound ACE2 | | | | | |
|---|---|---|---|---|---|---|---|---|---|---|---|
| Residue[a] | $\triangle E_{vdW}$ | $\triangle E_{elec}$ | $\triangle G_{polar}$ | $\triangle G_{non\text{-}polar}$ | $\triangle G_{binding}$ | Residue[a] | $\triangle E_{vdW}$ | $\triangle E_{elec}$ | $\triangle G_{polar}$ | $\triangle G_{non\text{-}polar}$ | $\triangle G_{binding}$ |
| S19 | -2.6 | 34.3 | 18.0 | -0.7 | 49.0 | S19 | -2.8 | 51.1 | 18.6 | -1.0 | 66.0 |
| Q24 | -12.3 | -7.4 | 11.2 | -1.4 | -9.9 | Q24 | -11.9 | -12.7 | 11.8 | -1.3 | -14.0 |
| T27 | -12.4 | -3.0 | 5.5 | -1.8 | -11.6 | T27 | -11.6 | -6.3 | 5.9 | -1.7 | -13.7 |
| F28 | -5.5 | -0.1 | -0.1 | -0.4 | -6.1 | F28 | -6.0 | -3.3 | 1.1 | -0.4 | -8.7 |
| D30 | -4.1 | -66.5 | 6.9 | -0.3 | -64.0 | D30 | -5.2 | -226.5 | 77.1 | -1.1 | -155.7 |
| K31 | -11.8 | 56.6 | 16.3 | -1.6 | 59.5 | K31 | -13.6 | -10.6 | 42.7 | -2.2 | 16.3 |
| H34 | -13.4 | 0.5 | 7.8 | -1.9 | -7.0 | H34 | -14.6 | -11.7 | 7.4 | -2.0 | -20.9 |
| E35 | -3.2 | -41.1 | -1.9 | -0.2 | -46.5 | E35 | -4.2 | -64.2 | 9.8 | -0.4 | -59.0 |
| E37 | -2.5 | -90.6 | 13.1 | -0.3 | -80.3 | E37 | -2.9 | -153.9 | 33.4 | -0.5 | -124.0 |
| D38 | -3.5 | -63.9 | 18.5 | -0.7 | -49.7 | D38 | -7.7 | -112.5 | 27.1 | -1.0 | -94.1 |
| Y41 | -11.7 | -7.8 | 5.0 | -1.0 | -15.5 | Y41 | -10.3 | -3.7 | 3.6 | -1.0 | -11.4 |
| Q42 | -5.2 | -2.2 | 1.9 | -0.7 | -6.2 | Q42 | -3.5 | -8.4 | 4.6 | -0.6 | -8.0 |
| L45 | -5.4 | 0.1 | 0.3 | -0.7 | -5.7 | L45 | -2.9 | 2.6 | -0.4 | -0.3 | -1.0 |
| L79 | -2.4 | 0.8 | 0.2 | -0.5 | -1.8 | L79 | -4.6 | -1.4 | 1.4 | -0.6 | -5.2 |
| M82 | -3.9 | -0.7 | 1.1 | -0.9 | -4.5 | M82 | -5.5 | -0.9 | 1.3 | -0.8 | -5.8 |
| Y83 | -1.1 | -17.5 | 5.3 | -0.4 | -13.7 | Y83 | -6.0 | -14.8 | 7.4 | -0.9 | -14.4 |
| E329 | 1.1 | -225.7 | 66.7 | -0.9 | -158.7 | E329 | -0.9 | -83.2 | 0.8 | 0.0 | -83.3 |
| N330 | -5.0 | -4.7 | 6.9 | -0.7 | -3.5 | N330 | -3.7 | -1.4 | 4.6 | -0.7 | -1.2 |
| K353 | -21.6 | 9.4 | 57.8 | -2.7 | 43.0 | K353 | -20.1 | 46.6 | 44.2 | -2.6 | 68.2 |
| G354 | -6.9 | -5.0 | 4.0 | -0.7 | -8.6 | G354 | -7.0 | -0.4 | 2.5 | -0.7 | -5.6 |
| D355 | -8.2 | -119.1 | 16.6 | -0.3 | -111.0 | D355 | -7.6 | -117.9 | 16.8 | -0.4 | -109.2 |
| R357 | -3.2 | 98.9 | -1.6 | -0.3 | 93.8 | R357 | -2.7 | 96.4 | -2.0 | -0.3 | 91.4 |
| R393 | -1.1 | 75.3 | -1.6 | -0.1 | 72.4 | R393 | -1.9 | 110.3 | -7.8 | -0.2 | 100.4 |
| **Sum1** | -130.7 | -122.3 | 187.4 | -16.8 | -82.4 | **Sum1** | -137.4 | -264.5 | 230.9 | -17.9 | -188.9 |
| **Sum2** | -4.3 | -225.6 | 67.1 | -1.6 | -164.4 | **Sum2** | -15.8 | -181.9 | 80.4 | -2.3 | -119.6 |
| **Sum3** | -135.0 | -347.9 | 254.5 | -18.4 | -246.8 | **Sum3** | -153.2 | -446.4 | 311.3 | -20.2 | -308.5 |

[a] ACE2 interface residues (i.e., IRCN-forming residues from ACE2) are colored the same as in Figures 5A and B in the main body of the paper, with the shared and non-shared ACE2 residues between IRCNs of the two complexes colored green and light green, respectively; the non-interface ACE2 residues (i.e., those do not make close contact with RBDs) are colored black; Sum1-3 indicate the cumulative values of respective energy terms for the shared, non-shared, and all ACE2 interface residues, respectively.

**Table S4.** Values of the residue binding free energy and energy components (kJ/mol) for RBM residues from $RBD_{CoV}$ and $RBD_{CoV2}$.

| $RBD_{CoV}$ | | | | | | $RBD_{CoV2}$ | | | | | |
|---|---|---|---|---|---|---|---|---|---|---|---|
| Residue[a] | $\triangle E_{vdW}$ | $\triangle E_{elec}$ | $\triangle G_{polar}$ | $\triangle G_{non\text{-}polar}$ | $\triangle G_{binding}$ | Residue[a] | $\triangle E_{vdW}$ | $\triangle E_{elec}$ | $\triangle G_{polar}$ | $\triangle G_{non\text{-}polar}$ | $\triangle G_{binding}$ |
| T438 | -0.1 | 0.5 | 0.1 | 0.0 | 0.5 | S438 | -0.1 | 0.8 | 0.0 | 0.0 | 0.7 |
| R439 | 1.5 | -619.8 | 67.9 | -1.0 | -551.4 | N439 | -0.4 | -3.6 | -0.4 | 0.0 | -4.4 |
| N440 | -0.2 | 2.0 | -0.3 | 0.0 | 1.6 | N440 | -0.1 | -1.6 | 0.0 | 0.0 | -1.7 |
| I441 | -0.1 | -0.7 | 0.3 | 0.0 | -0.5 | L441 | -0.1 | -0.2 | 0.1 | 0.0 | -0.2 |
| D442 | -0.1 | 387.6 | -4.8 | 0.0 | 382.7 | D442 | -0.1 | 373.4 | -3.5 | 0.0 | 369.8 |
| A443 | -0.2 | -5.4 | 0.6 | 0.0 | -5.0 | S443 | -0.3 | -6.7 | 0.5 | 0.0 | -6.5 |
| T444 | -0.4 | 7.5 | -1.3 | 0.0 | 5.8 | K444 | -0.4 | -381.5 | 1.6 | 0.0 | -380.3 |
| S445 | -0.9 | 3.2 | 0.1 | -0.1 | 2.3 | V445 | -0.7 | -6.1 | 0.1 | 0.0 | -6.8 |
| T446 | -1.0 | 2.3 | -0.3 | 0.0 | 1.0 | G446 | -1.8 | -0.5 | 0.8 | -0.2 | -1.6 |
| G447 | -0.7 | -3.5 | 0.6 | 0.0 | -3.6 | G447 | -0.9 | -3.7 | 0.6 | 0.0 | -3.9 |
| N448 | -0.4 | 2.3 | 0.2 | 0.0 | 2.0 | N448 | -0.5 | 3.7 | -0.4 | 0.0 | 2.9 |
| Y449 | -3.1 | -34.2 | 10.5 | -1.1 | -27.8 | Y449 | -5.1 | 1.9 | 3.0 | -0.8 | -1.0 |
| N450 | -0.1 | -3.3 | 0.2 | 0.0 | -3.1 | N450 | -0.1 | -4.8 | 0.2 | 0.0 | -4.6 |
| Y451 | -0.2 | 4.0 | -0.1 | 0.0 | 3.7 | Y451 | -0.2 | 4.7 | -0.4 | 0.0 | 4.0 |
| K452 | -0.3 | -416.8 | 4.8 | 0.0 | -412.3 | L452 | -0.3 | -3.5 | 0.2 | 0.0 | -3.5 |
| Y453 | -3.2 | 2.7 | 1.9 | -0.4 | 1.0 | Y453 | -3.4 | -1.9 | 4.6 | -0.5 | -1.2 |
| R454 | -0.3 | -351.7 | 2.1 | 0.0 | -349.9 | R454 | -0.4 | -342.2 | 3.5 | 0.0 | -339.1 |
| Y455 | -12.0 | 3.4 | 5.6 | -1.5 | -4.6 | L455 | -10.6 | 7.8 | -1.7 | -0.9 | -5.5 |
| L456 | -4.3 | -2.1 | 0.2 | -0.5 | -6.8 | F456 | -10.1 | -2.8 | 2.4 | -1.2 | -11.7 |
| R457 | -0.2 | -327.0 | 1.4 | 0.0 | -325.9 | R457 | -0.3 | -317.3 | 2.7 | 0.0 | -314.9 |
| H458 | -0.3 | -0.4 | 0.0 | 0.0 | -0.7 | K458 | -0.4 | -319.3 | 2.8 | 0.0 | -317.0 |
| G459 | 0.0 | 1.8 | -0.1 | 0.0 | 1.6 | S459 | -0.1 | 3.2 | -0.3 | 0.0 | 2.8 |
| K460 | -0.1 | -361.4 | 0.9 | 0.0 | -360.6 | N460 | -0.1 | -1.7 | 0.1 | 0.0 | -1.7 |
| L461 | 0.0 | 1.0 | 0.0 | 0.0 | 1.0 | L461 | 0.0 | 1.3 | 0.0 | 0.0 | 1.3 |
| R462 | 0.0 | -287.0 | 0.3 | 0.0 | -286.7 | K462 | 0.0 | -279.6 | 0.6 | 0.0 | -279.0 |
| P463 | 0.0 | -2.6 | 0.1 | 0.0 | -2.6 | P463 | 0.0 | -2.8 | 0.0 | 0.0 | -2.7 |
| F464 | 0.0 | -3.2 | 0.1 | 0.0 | -3.1 | F464 | 0.0 | -3.6 | 0.2 | 0.0 | -3.5 |
| E465 | 0.0 | 303.2 | -0.8 | 0.0 | 302.4 | E465 | 0.0 | 294.7 | -1.5 | 0.0 | 293.2 |
| R466 | 0.0 | -310.0 | 1.2 | 0.0 | -308.9 | R466 | 0.0 | -299.9 | 1.8 | 0.0 | -298.1 |
| D467 | 0.0 | 327.4 | -1.4 | 0.0 | 325.9 | D467 | 0.0 | 320.5 | -2.8 | 0.0 | 317.7 |
| I468 | 0.0 | 0.9 | 0.0 | 0.0 | 0.8 | I468 | 0.0 | 1.6 | 0.0 | 0.0 | 1.6 |
| S469 | 0.0 | 3.8 | -0.1 | 0.0 | 3.7 | S469 | 0.0 | 6.1 | -0.2 | 0.0 | 5.9 |
| N470 | -0.1 | 2.3 | 0.0 | 0.0 | 2.3 | T470 | -0.1 | 0.1 | 0.1 | 0.0 | 0.2 |
| V471 | -0.1 | 3.2 | -0.1 | 0.0 | 3.0 | E471 | -0.1 | 310.9 | -0.7 | 0.0 | 310.2 |
| P472 | -0.2 | -4.8 | 0.0 | 0.0 | -5.0 | I472 | -0.2 | -4.2 | 0.0 | 0.0 | -4.5 |
| F473 | -2.3 | 0.9 | 0.7 | -0.1 | -0.8 | Y473 | -2.6 | -3.1 | 2.6 | -0.2 | -3.3 |
| S474 | -1.1 | -1.4 | 0.2 | 0.0 | -2.2 | Q474 | -1.0 | -1.1 | -1.2 | 0.0 | -3.3 |
| P475 | -11.4 | -2.4 | 2.9 | -1.6 | -12.5 | A475 | -5.2 | -19.7 | 11.6 | -1.2 | -14.5 |
| D476 | -3.0 | 318.0 | 16.3 | -0.7 | 330.6 | G476 | -4.3 | -4.7 | 2.1 | -0.4 | -7.3 |
| G477 | -0.3 | -4.2 | 0.1 | 0.0 | -4.3 | S477 | -2.5 | -9.5 | 3.5 | -0.5 | -9.0 |
| K478 | -0.5 | -300.8 | -3.7 | -0.1 | -305.1 | T478 | -0.6 | 1.1 | 0.5 | 0.0 | 1.0 |
| P479 | -0.1 | 1.8 | 0.2 | 0.0 | 1.9 | P479 | -0.1 | 4.7 | 0.1 | 0.0 | 4.7 |
| C480 | -0.1 | -0.9 | 0.0 | 0.0 | -1.0 | C480 | -0.2 | 3.3 | 0.0 | 0.0 | 3.1 |
| T481 | -0.1 | 0.5 | 0.0 | 0.0 | 0.5 | N481 | -0.1 | 0.4 | 0.0 | 0.0 | 0.3 |
| - | - | - | - | - | - | G482 | 0.0 | 0.0 | 0.0 | 0.0 | -0.1 |
| P483 | -0.1 | -6.1 | 0.0 | 0.0 | -6.3 | V483 | -0.1 | -1.8 | -0.1 | 0.0 | -2.1 |
| P484 | -0.3 | -6.1 | 0.0 | 0.0 | -6.3 | E484 | -0.9 | 378.2 | 10.2 | -0.2 | 387.3 |
| A485 | -0.5 | 2.7 | 0.2 | 0.0 | 2.5 | G485 | -1.2 | -1.5 | 1.1 | -0.1 | -1.6 |
| L486 | -5.4 | -8.6 | 1.2 | -1.2 | -14.0 | F486 | -15.6 | -11.7 | 5.3 | -2.2 | -24.2 |

| | | | | | | | | | | | |
|---|---|---|---|---|---|---|---|---|---|---|---|
| N487 | -9.3 | -15.5 | 14.8 | -1.7 | -11.7 | N487 | -7.6 | -20.6 | 14.1 | -0.9 | -15.1 |
| C488 | -1.2 | 1.1 | 0.5 | 0.0 | 0.4 | C488 | -1.0 | 1.6 | 0.4 | 0.0 | 1.0 |
| Y489 | -16.8 | -5.0 | 7.5 | -2.0 | -16.2 | Y489 | -14.7 | 0.8 | 8.0 | -1.8 | -7.7 |
| W490 | -1.3 | 2.7 | 0.7 | -0.1 | 2.0 | F490 | -1.2 | 2.9 | 2.0 | -0.1 | 3.5 |
| P491 | -0.5 | 4.3 | -0.3 | 0.0 | 3.5 | P491 | -0.5 | 5.9 | -0.9 | 0.0 | 4.6 |
| L492 | -0.7 | 2.5 | 0.7 | 0.0 | 2.5 | L492 | -0.7 | -2.6 | 2.9 | -0.1 | -0.5 |
| N493 | -7.1 | -11.3 | 9.5 | -1.0 | -9.9 | Q493 | -7.5 | -61.2 | 23.9 | -1.7 | -46.6 |
| D494 | -2.7 | 501.0 | -9.4 | -0.2 | 488.7 | S494 | -1.7 | 13.4 | -0.9 | -0.1 | 10.8 |
| Y495 | -2.1 | -13.2 | 6.4 | -0.3 | -9.2 | Y495 | -1.8 | -7.3 | 0.8 | 0.0 | -8.3 |
| G496 | -3.0 | -10.0 | 5.5 | -0.2 | -7.7 | G496 | -1.7 | -19.9 | 6.6 | -0.6 | -15.6 |
| F497 | -1.7 | -2.5 | 1.2 | 0.0 | -3.0 | F497 | -2.1 | 0.1 | -0.2 | 0.0 | -2.2 |
| Y498 | -14.6 | -0.7 | 4.2 | -1.6 | -12.6 | Q498 | -8.1 | -34.1 | 13.2 | -1.3 | -30.3 |
| T499 | -1.8 | 6.9 | -3.3 | 0.0 | 1.8 | P499 | -1.8 | 3.1 | -0.1 | 0.0 | 1.2 |
| T500 | -13.8 | -17.8 | 14.5 | -2.1 | -19.3 | T500 | -12.9 | -13.1 | 11.7 | -2.3 | -16.6 |
| T501 | -14.0 | -12.5 | 5.3 | -0.9 | -22.1 | N501 | -15.2 | -24.4 | 14.5 | -0.8 | -25.9 |
| G502 | -6.5 | -9.1 | 4.0 | -1.0 | -12.6 | G502 | -5.1 | -10.0 | 3.5 | -0.8 | -12.3 |
| I503 | -3.6 | -2.8 | 0.0 | -0.5 | -7.0 | V503 | -2.0 | -2.7 | -0.4 | -0.1 | -5.3 |
| G504 | -0.9 | -7.1 | 1.1 | 0.0 | -7.0 | G504 | -0.9 | -4.9 | 0.5 | 0.0 | -5.3 |
| Y505 | -17.1 | -13.8 | 7.3 | -1.7 | -25.3 | Y505 | -18.5 | -31.7 | 12.9 | -2.3 | -39.6 |
| Q506 | -1.7 | -16.8 | 3.6 | -0.1 | -14.9 | Q506 | -1.4 | -11.7 | 1.3 | 0.0 | -11.8 |
| **Sum** | -172.9 | -1300.8 | 181.9 | -21.8 | -1313.7 | **Sum** | -177.6 | -538.7 | 163.3 | -21.3 | -574.3 |

[a] RBM spans residues 438-506 in the two RBDs; the residue numbering is according to $RBD_{CoV2}$. Sum indicates the cumulative values of respective energy terms for the RBM residues.
